# Ras Activation by Hydrostatic Pressure is Enhanced by GAP and GEF *in vitro*

**DOI:** 10.1101/2023.12.05.570111

**Authors:** Teruhiko Matsuda, Minki Chang, Katsuko Furukawa, Takashi Ushida, Taro Q.P. Uyeda

## Abstract

Hydrostatic pressure (HP) is a necessary stimulus for cell differentiation and growth in cultured chondrocytes. Assuming that Ras activation is involved in HP-induced reactions, if cellular Ras activity is increased by HP, Ras itself, Raf, Ras regulators including GTPase activating proteins (GAP) and guanine nucleotide exchange factors (GEF), then the upstream receptor and/or interactions between them should have HP sensitivity. Based on this hypothesis, we first attempted to examine whether Ras is activated by HP, and in the affirmative, identify which factors displayed HP sensitivity using an *in vitro* system to measure Ras activity. This *in vitro* system included mRaichu, a FRET-based Ras activity probe, to which two-point mutations were introduced to reduce Ras-independent signals. This improved Raichu was used to investigate the HP sensitivity of two components, the GAP domain (GAPd) derived from p120GAP and the GEF domain (GEFd) derived from hSOS-1. It was found that HP weakly activated Ras activity in the absence of GAPd and GEFd, presumably by facilitating GDP dissociation from Ras. A low concentration of GAPd enhanced HP-induced Ras activation by 16.3% whereas high concentrations of GAPd removed HP sensitivity, suggesting that HP partially dissociates GAPd from the GAPd-Ras-GDP complex and reduces the fraction of inactive Ras. Moreover, a broad concentration range of GEFd also enhanced HP-induced Ras activation. Given that HP also increased Ras activity under a condition mimicking cellular Ras activity, we propose that Ras activation is involved in the differentiation and growth stimulation of chondrocytes by HP.

**Statement of Significance:** The Ras-cycle has been implicated in the regulation of growth and differentiation of eukaryotic cells. Here, we investigated the relationship between hydrostatic pressure (HP) and components of the Ras-cycle: Ras-Raf, the GAP domain, and the GEF domain. Generally, HP tends to weaken protein-protein and protein-ligand interactions, but in this study, a seemingly positive response was observed: HP-induced Ras activation. Additionally, this response was enhanced by the GEF domain and the GAP domain. Thus, given the multiple cellular functions of Ras and the responses of the Ras-cycle to HP, this study will help clarify the molecular mechanism by which HP modulates cellular functions, particularly in chondrocytes, which are subjected to repetitive HP stimuli *in vivo*.

## Introduction

Changes in the external environment, including hydrostatic pressure (HP), induce various cellular responses (reviewed by (1,2)). Chondrocytes in joints are subjected to repetitive HP stimuli, and HP induces not only cartilage deformation but also regulates chondrocyte metabolic activity. Additionally, the growth of chondrocytes in culture is enhanced by moderate HP stimulation (0.5-5 MPa)^(3–5)^. In contrast, 25 MPa of HP, which is the higher end of the physiological HP range, increases the expression of stress-related and apoptosis-related genes but decreases the expression of cartilage matrix genes, leading to osteoarthritis in joints^(6)^. Cellular HP responses are closely related to the HP sensitivity of proteins themselves. Recent studies showed that HP induces structural changes in specific proteins such as F1 ATPase^(7)^, microtubules^(8,9)^, Staphylococcal nuclease (SNase)^(10)^ and ubiquitin^(11)^. In our own attempt to characterize the HP-induced response of chondrocytes, preliminary results suggested that HP stimulation (10 MPa) induces the activation of intracellular Ras in cultured mouse chondrocytes (ATDC5) (T. Ushida, unpublished data). Ras is one of several small GTPases implicated in the regulation of the signal transduction system for proliferation, differentiation (reviewed by (12,13)), cell motility^(14)^, axon formation^(15–17)^, cell adhesion^(18)^, and gene expression underlying those events (reviewed by (19)). Ras assumes two states, the “active” state when it binds GTP, and the “inactive” state when it binds GDP (reviewed by (13,20–23)). These two states are interchangeable, forming the Ras-cycle. Active Ras binds specifically to the Ras binding domain of Raf (RafRBD)^(24)^ and this complex is implicated in the regulation of cell growth and differentiation by modulating the expression of various genes^(25,26)^. Moreover, many cancer cells are characterized by unregulated Ras activation (reviewed by (27)). In the cell, the transitions between active and inactive Ras are accelerated by GTPase activating proteins (GAPs) and guanine nucleotide exchange factors (GEFs) (reviewed by (23,28)). These regulators adjust the ratio of active and inactive Ras and regulate Ras-related phenomena and carcinogenesis (reviewed by (13,20–22,28)). Based on the Ras-cycle framework, the HP response observed in cultured mouse chondrocytes should derive from HP-sensitivity of Ras itself, Ras regulators (GAPs, GEFs), interactions between them, as well as the interaction between Ras and RafRBD and/or the upstream growth factor receptor. However, because HP generally tends to dissociate protein oligomers and weakens protein-protein interactions (reviewed by (29)), the upstream receptor (receptor tyrosine kinases) that dimerizes upon activation (reviewed by (30)) is unlikely to confer the positive HP sensitivity of Ras in chondrocytes. Regarding the HP sensitivity of Ras itself, NMR spectroscopy suggested that the GTP-bound state of Ras is divided into three sub-states: the GEF interacting state, the Raf interacting state and the GAP interacting state; moreover, as HP was increased to 20-150 MPa, the relative populations of the first states decreased while that of the last two state increased^(31)^. Additionally, that NMR study suggested that observed pressure-sensitive regions of Ras are closely located at the interaction sites with effectors, including Raf, and regulators such as GEFs and GAPs. Hence, it is hypothesized that the HP sensitivity of the Ras-cycle resides in Ras itself, Ras regulators, interactions between them, and/or the interaction between Ras and RafRBD. Identifying the major HP-sensitive process among these mechanisms is difficult using an *in vivo* system, however. Therefore, we set out to develop an *in vitro* system to measure HP-induced time-dependent changes of Ras activity using a Förster resonance energy transfer (FRET)-based probe for active Ras, Raichu, previously constructed by Mochizuki et al. ^(32)^. Raichu is a chimeric protein consisting of four domains, YFP, H-Ras (hereafter referred to as Ras), RafRBD, and CFP, in this order. Raichu can assume two distinct forms, open and closed, corresponding to the state of Ras, owing to the specific intramolecular binding between active Ras and RafRBD. In the closed form, CFP and YFP moieties come close to each other, allowing FRET to occur so that YFP fluorescence is observed when CFP is excited. In contrast, when Ras is in the inactive state, Raichu supposedly takes the open form, so that FRET efficiency decreases and CFP fluorescence is emitted when CFP is excited. By using this probe, Ras activity was visualized in a living cell using a fluorescence microscope^(32)^. In our experiments using a fluorescence spectrophotometer, however, we noted that a certain fraction of Raichu molecules spontaneously form the closed configuration, even in the presence of GDP and in the absence of GTP. We reasoned that this erroneous closed configuration is due to the affinity between CFP and YFP moieties since CFP and YFP are variants of GFP and because it has been firmly established that GFP molecules tend to dimerize^(33)^. To minimize this artifactual signal, we introduced point mutations equivalent to A206K^(34,35)^ of GFP into CFP and YFP moieties within Raichu, to yield an improved probe named mRaichu. Using mRaichu, the Ras-GAP domain (GAPd) of p120-GAP (reviewed by (36)), the Ras-GEF domain (Cdc25-Rem domains) (GEFd) of hSOS-1^(37)^, an HP-resistant optical cell and a fluorescence spectrophotometer, we established an *in vitro* real-time Ras activity measurement system.

Using the established system, we detected HP-induced Ras activation, which is consistent with our preliminary result that HP induces Ras activation in mouse chondrocytes. Moreover, we measured Ras activity in a number of artificial and cell mimic conditions under the higher end (20–30 MPa) of the physiological HP range (0-20 MPa) on human chondrocytes in joints^(38–40)^ in order to elucidate the mechanism of HP sensitivity of the Ras-cycle. Our data indicate that Ras itself has weak HP sensitivity although the interaction between Ras and GAP and GEF is mainly responsible for the sensitivity of the Ras-cycle to HP. Our results are also consistent with the idea that cell behaviors that depend on Ras activity, including cell growth and differentiation, are modulated by HP in cells.

## Materials and Methods

### Construction of the expression plasmid and purification of mRaichu

mRaichu is a derivative of Raichu^(32)^, which has additional point mutations in YFP and CFP moieties that are equivalent to A206K of GFP^(34,35)^. The mRaichu gene was constructed based on a PCR-based method and fused to the 3′ end of the His-tag coding sequence and the TEV recognition sequence. The His-TEV-mRaichu coding sequence was amplified by PCR using the primers GAC<u>GAATTC</u>ATGAATCACAAAGTGCATCAT and CTCGAC<u>AAGCTT</u>TTAGATTCTGTGCTTTTAAGC and was inserted into the *Sma*I site of pBCKS (Stratagene). The coding sequence in the resultant plasmid was subcloned into the pFastBac™ Dual Expression Vector (Thermo Fisher) using *Eco*RI and *Hind*III sites (underlined in the primer sequences). The coding sequence in this recombinant donor plasmid was inserted into the bacmid in *Escherichia coli* DH10Bac by homologous recombination. Thereafter, His-TEV-mRaichu was expressed in Sf9 insect cells, referring to the Bac-to-Bac^®^ Baculovirus Expression System user manual Version F (Invitrogen, 2010).

mRaichu was purified using a method developed for Ras purification^(41)^. All procedures were performed at 0-5 °C or on ice. To stabilize the structure of Ras in mRaichu, all buffers after the lysis step of this method contained 30 μM GDP and 1 mM MgCl_2_. Pelleted cells, after washing with buffer (10 mM Hepes pH 7.4, 100 mM NaCl), were lysed by resuspending them at 0.1 g cell/mL with mRaichu-Lysis buffer (20 mM Hepes pH 7.4, 100 mM NaCl, 7 mM 2-mercaptoethanol (βmE), 4 mM MgCl_2_, 30 μM GDP, 10 mM imidazole (pH 7.4), 5% glycerol, 10 µM phenyl-methyl sulfonyl fluoride (PMSF), 0.1% Triton-X 100, 10 μM leupeptin) and incubated for 30 min on ice. Soluble proteins from lysed insect cells were obtained by centrifugation at 10,000 rpm for 30 min at 4 °C, mixed with 1 mL Ni-NTA Superflow resin (GE-Healthcare) that had been equilibrated with mRaichu-Wash buffer 1 (20 mM Tris pH 7.4, 1 M NaCl, 7 mM βmE, 4 mM MgCl_2_, 30 μM GDP, 25 mM imidazole, 5% glycerol). The mixture was gently agitated for 1 h at 5 °C. After washing the Ni-NTA resin with 20 mL mRaichu-Wash buffer 1 and 10 mL mRaichu-Wash buffer 2 (20 mM Tris pH 7.4, 300 mM NaCl, 7 mM βmE, 4 mM MgCl_2_, 30 μM GDP, 25 mM imidazole, 5% glycerol), the target protein was eluted with 15 mL mRaichu-Elution buffer (20 mM Tris pH 7.4, 300 mM NaCl, 7 mM βmE, 4 mM MgCl_2_, 30 μM GDP, 300 mM imidazole, 5% glycerol). The eluate was dialyzed against 1 L mRaichu-Dialysis buffer (10 mM Tris pH 7.4, 50 mM NaCl, 7 mM βmE, 2 mM MgCl_2_, 30 μM GDP, 0.05% NaN_3_), concentrated using Amicon^®^ Ultra Centrifugal Filters (Millipore), with a cut-off molecular weight of 100 kDa, and clarified by ultracentrifugation at 80,000 rpm for 10 min. After calculating the protein concentration from the molar absorption coefficient, which was calculated from the amino acid sequence of mRaichu on PlotParam (Expasy) and absorption at 280 nm measured by Nanodrop 2000 (Thermo Fisher), the solution was supplemented with glycerol to 10% and rapidly frozen in small aliquots and stored at -80 °C. His-TEV-mRaichu purified using this method, hereafter simply called mRaichu, was used in this study without removing the His-tag.

### Construction of expression plasmids and purification of the Ras-GAP domain (GAPd) and the Ras-GEF domain (Rem-Cdc25 domains) (GEFd)

The GAPd coding sequence derived from the p120GAP coding sequence (*RASA1*; Accession: NP_002881) and the GEFd coding sequence derived from the h-SOS1 coding sequence (*SOS1*; Accession: SPT35795) in the cDNA library of HeLa cells were amplified by PCR. The sequences of the primers were CATCAT<u>GGATCC</u>CAGATGAGGCTGCCTA and ATGATG<u>GTCGAC</u>CTAGGTACCTGGTCTTGGGT for GAPd, and CATCAT<u>GGATCC</u>GAAAAAATCATGCCAGAAGAA and ATGATG<u>GTCGAC</u>CTACCTGACATCATTGGTTT for GEFd. The underlines show *Bam*HI and *Sal*I restriction sites added for subsequent subcloning. The PCR products were inserted into the *Sma*I site of the pBCKS plasmid, and the sequences of the PCR products were confirmed. Each coding sequence in pBCKS was subcloned into pCold-His-TEV^(42)^ using *Bam*HI and *Sal*I. His-TEV-GAPd and His-TEV-GEFd were expressed using each expression plasmid in *E. coli* BL21 (DE3, Takara).

His-TEV-GAPd and His-TEV-GEFd were purified as follows. All procedures were performed at 0-5 °C or on ice. Pelleted cells, after washing with 10 mM Hepes (pH 7.4) and 100 mM NaCl, were resuspended at 0.1 g cell/mL with lysis buffer (20 mM Hepes pH 7.4, 100 mM NaCl, 7 mM βmE, 10 mM imidazole, 10 µM PMSF, 0.1% Triton-X 100, 10 μM leupeptin). The cells were disrupted by sonication at maximum output 25% duty cycle for 8 min with 30 % of maximum amplitude using Digital Sonifier 250DA (BRANSON). Soluble proteins from the sonicated cells were obtained by centrifugation at 10,000 rpm for 30 min at 4 ℃ and His-TEV-GAPd and His-TEV-GEFd were purified in a manner similar to mRaichu using 0.5 mL Ni-NTA Superflow resin, 20 mL wash buffer 1 (20 mM Tris pH 7.4, 1 M NaCl, 7 mM βmE, 4 mM MgCl_2_, 25 mM imidazole), 10 mL wash buffer 2 (20 mM Tris pH 7.4, 300 mM NaCl, 7 mM βmE, 25 mM imidazole), 10 mL elution buffer (20 mM Tris pH 7.4, 300 mM NaCl, 7 mM βmE, 300 mM imidazole) and 500 mL dialysis buffer (10 mM Tris pH 7.4, 50 mM NaCl, 7 mM βmE, 0.05% NaN_3_) when these proteins were harvested from a 500 mL culture. The dialyzed solutions were then concentrated and clarified by ultracentrifugation at 80,000 rpm for 10 min. The clarified solutions were further purified by gel filtration chromatography (Superdex^TM^ 200 10/300 GL, Cytiva) for GEFd, or by ion exchange chromatography (Bio-scale^TM^ Mini Macro-Prep HighQ, BIORAD) for GAPd. The final products were stored at -80 ℃ after measuring the protein concentration as described for mRaichu above and supplementing with 10% glycerol. His-TEV-GAPd and His-TEV-GEFd purified using this method, simply referred to as GAPd and GEFd, respectively, were used without cleaving off the His-tag.

### Measurement of mRaichu FRET

To verify the function of mRaichu, the fluorescence spectra of mRaichu were measured in solutions containing 3 mg/mL BSA (SIGMA, Product No. 017-25771), 100 mM NaCl, 10 mM Hepes pH 7.4, 1 mM MgCl_2_, 10 mM dithiothreitol (DTT), 0.3 µM mRaichu and various additives including nucleotides, GAPd, GEFd and proteinase K. The mixtures were incubated for 1 h at 37 ℃ to reach a quasi-steady state and then introduced into a quartz cell (Hellma Analytics). Fluorescence spectra of the solution were measured using a fluorescence spectrophotometer A (FP-8500, Jasco) at an excitation wavelength of 433 nm. Although the temperature of the samples was not regulated when single measurements were made at ambient pressure, the measurements were made immediately after introducing the sample at 37 ℃ into the quartz cell before the temperature dropped to room temperature.

### Measurement of time-dependent FRET change of mRaichu under hydrostatic pressure and atmospheric pressure

Time-dependent changes in fluorescence spectra of mRaichu were measured by fluorescence spectrophotometry with and without HP. The solution conditions were identical between HP and atmospheric pressure (AP) measurements. The mixtures containing 0.3 µM mRaichu and various other components were incubated for 1 h at 37 ℃ to reach a quasi-steady state and then 300 µL of the solution was introduced for each measurement into the inner cell of the HP-resistant optical cell (pci-100, Syn Corporation) using a syringe. This inner cell was set in the outer HP-resistant optical cell. After filling the space between the inner and outer cells with deaerated water using a pump (Personal Pump NP-S-321, Nippon Precision Science), this HP-resistant cell assembly was set in a fluorescence spectrophotometer B (FP-6600, Jasco). Here, HP was indirectly applied to the solution in the inner cell through a flexible silicone portion of the inner cell’s surface. Fluorescence spectra were measured with an excitation light of 433 nm with and without approximately 25 MPa HP at 37 ℃.

### Calculation of the Ras activity ratio

Ras activity was calculated from the emission intensities of YFP and CFP when CFP was excited. Spectral measurements were made at 1 nm intervals using a 3 nm slit. Moreover, to improve the S/N ratio, when the time-dependent changes of the fluorescence intensity ratios were evaluated, they were calculated based on the summed fluorescence intensities over the range of 470-480 nm for CFP and 527-537 nm for YFP. Then, the 10 nm-averaged intensity of YFP was divided by that of CFP to yield the YFP/CFP ratio, and this value was then used as the indicator of Ras activity after correction of the bleed-through of CFP fluorescence in mRaichu to the YFP fluorescence window as follows. First, a proportionality constant of the bleed-through of CFP fluorescence in the YFP fluorescence window to the CFP fluorescence intensity in the CFP fluorescence window was determined from the spectrum data shown on the homepage of Thermo Fisher Scientific (https://www.thermofisher.com/order/fluorescence-spectraviewer/#!/). The observed YFP fluorescence intensity of mRaichu was corrected by subtracting the CFP fluorescence bleed-through calculated using this proportionality constant.

Moreover, as described in the Results, we noticed that the sensitivity of the two photodetectors of the two fluorescence spectrometers used in this study had different wavelength dependences, resulting in different YFP/CFP ratios, even when the same samples were measured. To allow a direct comparison of data obtained using the two fluorescence spectrometers, an additional correction was made as described in the Supporting Information.

The initial YFP/CFP ratio before HP was applied in the HP experiments and that in the AP experiments were sometimes unequal, even though those values should be identical under identical solution conditions. These differences presumably arose due to slight random variations in the incidence angles of excitation light. Therefore, the average of the initial values of all the HP and AP experiments, each of which consisted of multiple time-course measurements, was used as the initial YFP/CFP ratio for all the time-course measurements under that solution condition. All subsequent YFP/CFP ratios in each time-course measurement were translated such that the initial value of the measurement would be equal to the average of all the HP and AP experiments under that solution condition. This correction was made based on the assumption that the measurement-to-measurement variation of the initial value of each measurement remained unchanged throughout the measurement. Using those corrected YFP/CFP ratios, Ras activity [%] was calculated such that Ras activity [%] under the GDP condition was 0% and 100% in the presence of guanosine-5’-[(β,γ)-methyleno]triphosphate (GppCp). Ras activity [%] under AP was calculated from the YFP/CFP ratio of GDP and GppCp conditions under AP, whereas Ras activity [%] under HP was calculated from the same ratios under HP. Of note, despite the A206K-equivalent point mutations introduced into the CFP and YFP moieties, small but detectable fractions of mRaichu molecules were judged to spontaneously form the HP-sensitive closed configuration even under the GDP condition (Figure 3B), although the above calculation method cancelled the contributions of the spontaneous formation of closed configurations that were independent of Ras activities. To reduce the effect of outliers, the difference between Ras activity [%] under AP and HP at 5 min after the application of HP was defined as the average of the differences at 4 min and 6 min after HP was applied, due to increased variation in the YFP/CFP ratio under HP.

## Results

### Functionality of purified mRaichu and interaction with GAPd and GEFd

An *in vitro* system was constructed to measure Ras activity using Raichu (Figure 1A), a FRET-based Ras activity probe^(32)^. We first attempted to prepare Raichu using *E. coli*. However, most of the expressed Raichu passed into the insoluble fraction while refolding from the solubilized Raichu pellet was also unsuccessful. Therefore, soluble Raichu was prepared using an insect cell expression system (Figure S1A). When fluorescence spectra were observed using purified Raichu and excitation light for CFP, YFP fluorescence was significantly stronger in the presence of guanosine-5’-[(β,γ)-methyleno]triphosphate (GppCp), a non-hydrolyzable GTP analog, than in the presence of GDP, as expected (Figure S1B), leading us to conclude that the purified Raichu is a functional reporter of Ras activity. As shown in Figure S1C, HP induced a decrease of the YFP/CFP ratio in the presence of GDP or GppCp when time-dependent changes of CFP and YFP fluorescence intensities of Raichu were measured, while no significant changes were detected in the presence of proteinase K. The HP-induced change of the fluorescence spectrum of Raichu in the presence of GDP suggested that a certain fraction of Raichu molecules spontaneously formed the closed configuration in an HP-sensitive manner even when Ras was in an inactive state. This HP-sensitive closed state can be caused by the affinity between Ras-GDP and Raf-derived RBD moieties and/or CFP and YFP moieties within Raichu. Of these two possible mechanisms, we decided to deal with the latter because it is an artifact unrelated to the Ras cycle since the CFP-YFP interaction was plausible since CFP and YFP are variants of GFP, and because it is well established that GFP molecules tend to dimerize^(33)^. To minimize this artifactual signal, we took advantage of the A206K mutation of GFP, which was shown to decrease the affinity between two GFP molecules^(34,35)^. We introduced point mutations equivalent to A206K of GFP into CFP and YFP moieties, to yield mRaichu (Figure 1A). Figure 1B and C show the purification of mRaichu from insect cells. A protein with the expected molecular weight was expressed and approximately half of this protein was soluble (Figure 1B). Thus, this soluble protein was purified with Ni-resin. The purity of thus obtained mRaichu was 78-84%, as estimated by SDS-PAGE (Figure 1B). Aggregation or oligomerization was not detected in the purified mRaichu in the experimental solution containing 0.3 µM mRaichu and 100 µM GDP (pH 7.4) following examination by gel filtration chromatography (Figure 1C). We next characterized this protein in terms of FRET efficiency, which reflects the fraction of active Ras in mRaichu (Figure 1D, E). The mixture containing mRaichu and other components was brought to a quasi-static state by incubation for 1 h and then injected into the quartz cell and measured by fluorescence spectrophotometer A. Figure 1D shows the fluorescence spectra in the presence of 1 mM GDP, 1 mM GTP or 3 mM GppCp. We also made measurements in the presence of a mixture of 19.8 µM GDP and 13.2 µM GTP, in which the spectrum was halfway between that in the presence of 1 mM GDP and that in the presence of 3 mM GppCp (referred to as the DT condition hereafter). Where YFP and CFP moieties were considered completely independent due to the cleavage of the linker between Ras and Raf in mRaichu, effects of proteinase K were also examined in the DT condition. Figure 1E shows the fluorescence intensity of YFP divided by that of CFP (YFP/CFP ratio), which reflects the fraction of mRaichu molecules in which the Ras moiety is bound to the RafRBD moiety and hence is a quantitative indicator of the active fraction of Ras. The fluorescence intensity of YFP was higher than that of CFP under the GDP, GppCp, DT and GTP conditions (i.e., YFP/CFP ratio >1) although, as expected, the YFP/CFP ratios differed among the four conditions (Figure 1D, E). These spectral differences among the different conditions indicates that mRaichu changes its form, thus reflecting the activity of Ras (Figure 1A). The fluorescence spectra did not change when the concentration of GDP or GppCp was further increased from 1 mM, indicating that 1 mM of GDP and 3 mM GppCp were saturated (Figure S2). Furthermore, in the DT + proteinase K condition, the YFP/CFP ratio was lower than that of the GDP condition. This result supports our conclusion that the GDP- and GppCp-dependent spectral changes of mRaichu were caused by intramolecular FRET, as had been shown previously by Mochizuki et al. in *in vivo* experiments^(32)^.

**Figure 1.**
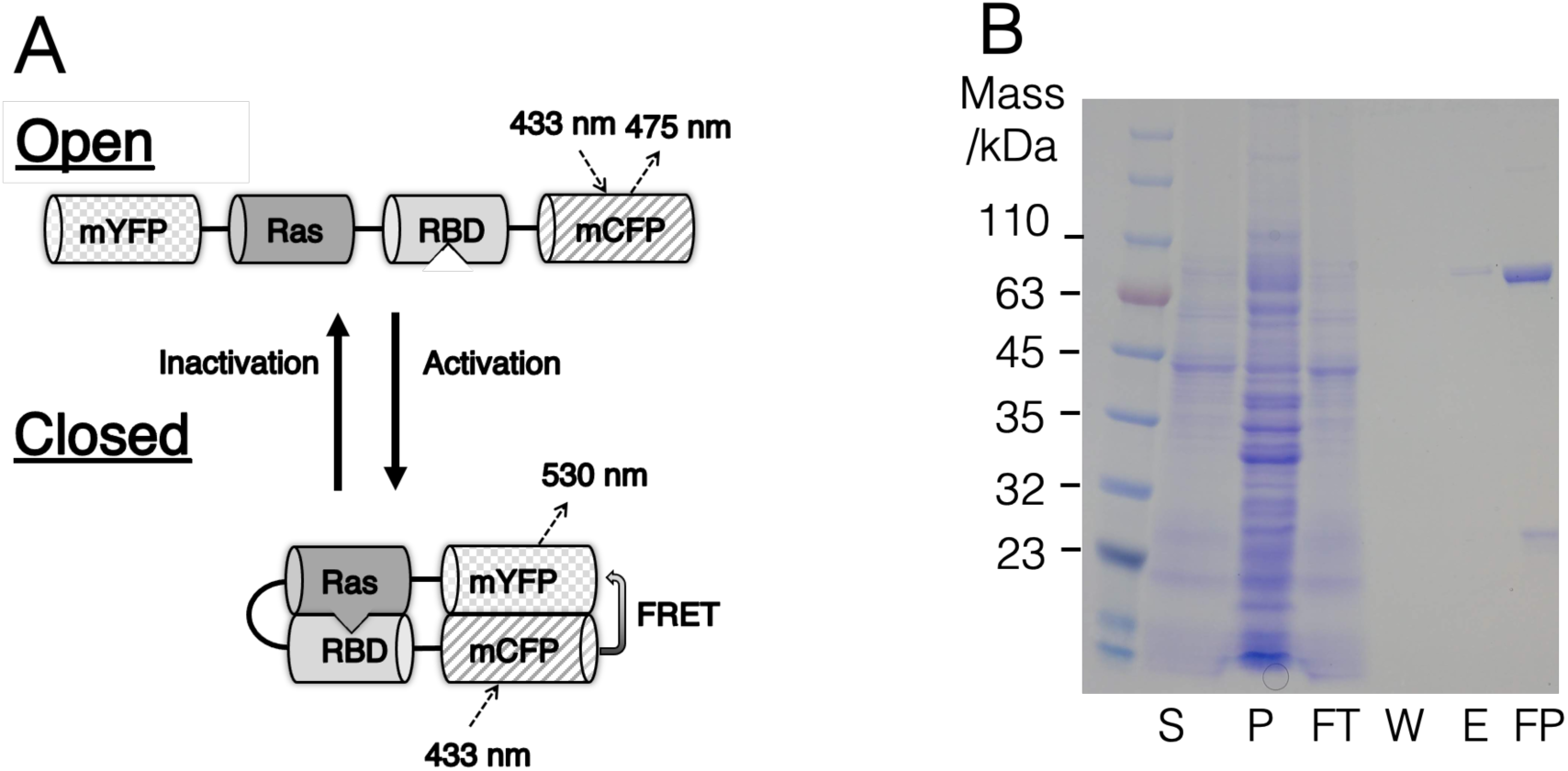
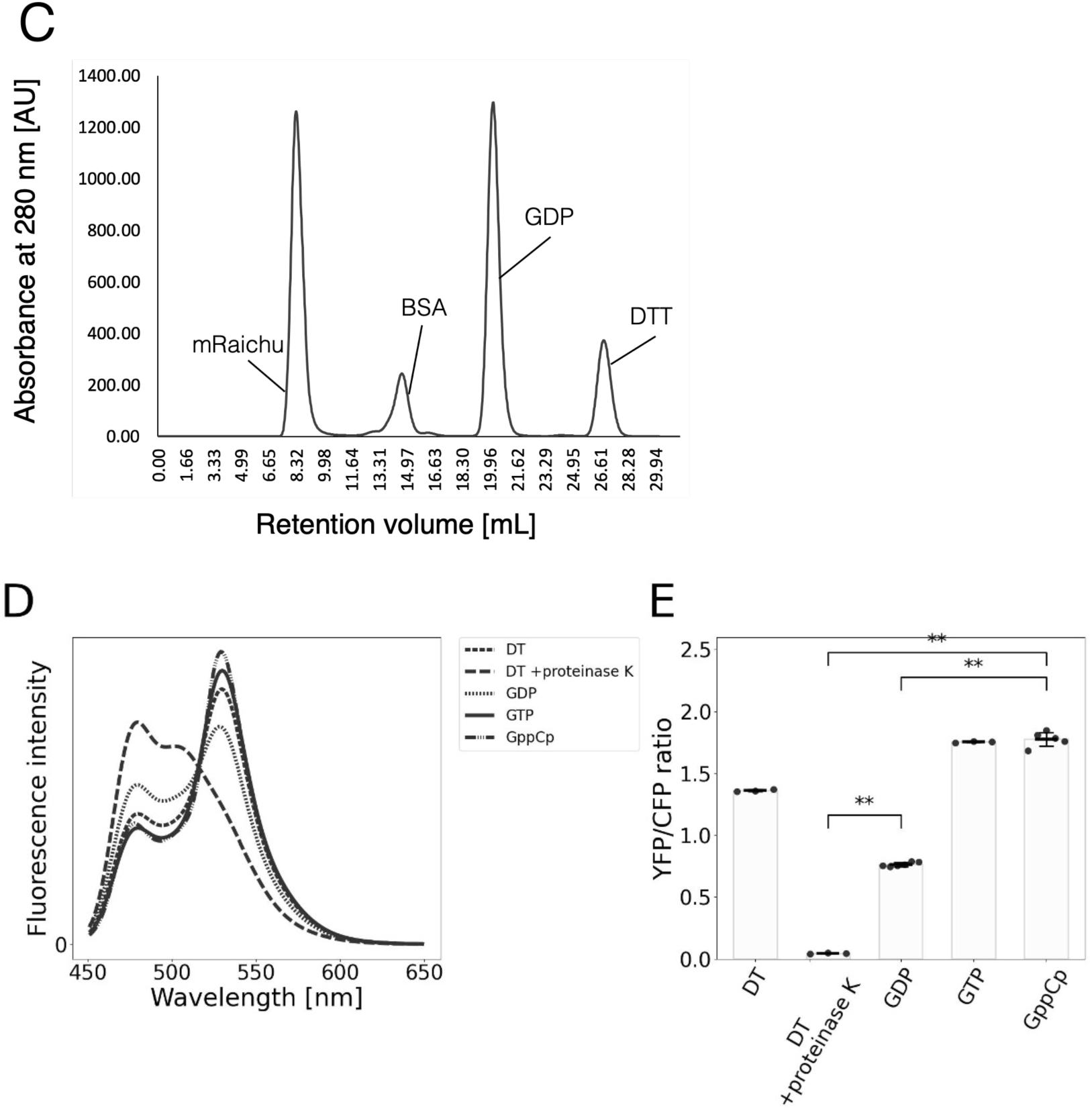
Purification of mRaichu (A) Schematic structure of mRaichu. mCFP and mYFP are CFP and YFP moieties carrying the A206K-equivalent mutations, respectively. (B) Purification of mRaichu (predicted molecular mass is 87,000). S, P: Supernatant and pellet fractions after insect cell lysis and centrifugation, FT: Flow-through after incubation with Ni-NTA resin, W: Wash effluent with Ni-NTA washing, E: Eluted solution, and FP: Final product after concentration and ultracentrifugation. (C) Gel filtration chromatography of the final product of mRaichu. BSA was included as the marker. (D) Difference in fluorescence spectra of mRaichu depending on the solution conditions (average of 6 measurements). Fluorescence spectra of the mixtures were measured at an excitation wavelength of 433 nm. (E) YFP/CFP ratio ± SD calculated from the data in (D). The horizontal black bars and error bars indicate the average value ± SD. GDP, GppCp and GTP conditions are the solutions containing 0.3 µM mRaichu and 1 mM GDP, 3 mM GppCp or 1 mM GTP, respectively. The DT condition contained 0.3 µM mRaichu, 19.8 µM GDP and 13.2 µM GTP. The DT + proteinase K condition contained 100 μg/mL proteinase K. **: the probability of no difference between Ras activities under HP and AP is lower than the significance level (*t* = 0.01; Welch’s *t*-test).

The Ras-GAP domain (GAPd) of human p120-GAP and the Ras-GEF domain (Cdc25 domain and Rem domain) (GEFd) of hSOS-1 were prepared to reconstitute the Ras-cycle. These proteins were expressed in *E. coli* and purified with Ni-resin (Figure S3A, B). Figure 2 shows the effects of these proteins on mRaichu. GAPd added to mRaichu in the DT condition reduced the YFP/CFP ratio in a concentration-dependent manner. This result demonstrated that GAPd increased the fraction of inactive Ras, as expected (Figure 2A, C). Conversely, GEFd added to mRaichu in the DT condition elevated the YFP/CFP ratio, demonstrating that GEFd increased the fraction of active Ras, again as expected (Figure 2B, C). Based on these results, we concluded that the prepared GAPd and GEFd were functional in terms of regulating Ras activity.

**Figure 2.**
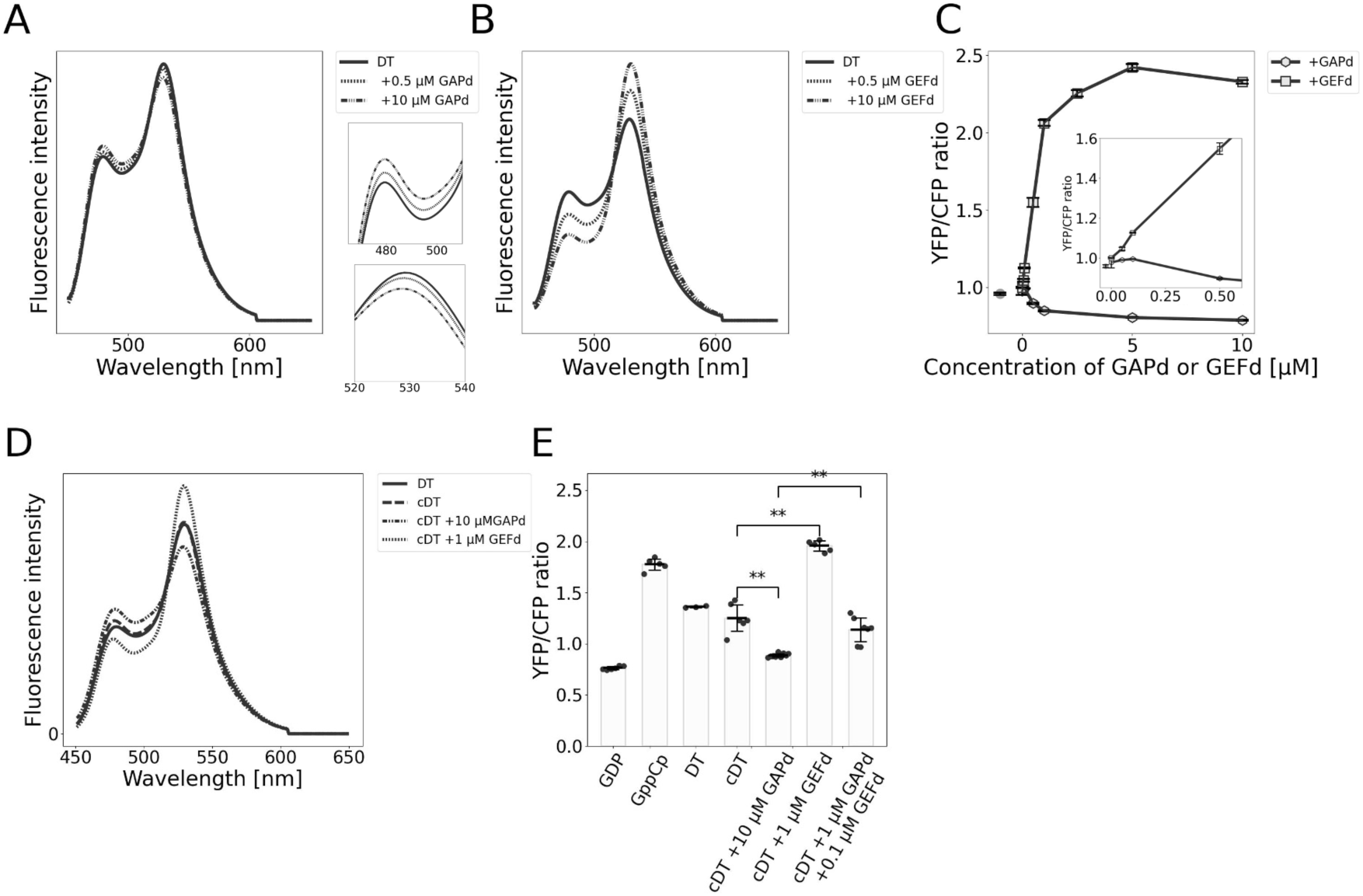
Effects of GAPd and GEFd on the fluorescence spectra of mRaichu. (A, B) Fluorescence spectra of mRaichu in the presence of various concentrations of GAPd and GEFd (average of 3 independent measurements). Excitation was at 433 nm. (C) The data were calculated from those shown in (A) ad (B). The gray data point near the vertical axis indicates the YFP/CFP ratio of the DT condition shown as the reference. The inset in (C) shows the enlarged view at low concentrations of GAPd and GEFd. (D) Fluorescence spectra of mRaichu in the presence of various concentrations of GAPd and GEFd (average of 6 measurements). Excitation was at 433 nm. (E) YFP/CFP ratio ± SD calculated from the YFP/CFP ratios of the data in the following solutions. The horizontal black bars and error bars indicate the average value ± SD. GDP and GppCp conditions contained 0.3 µM mRaichu and 1 mM GDP or 3 mM GppCp, respectively. The DT condition contained 0.3 µM mRaichu, 19.8 µM GDP and 13.2 µM GTP and the cDT condition contained 0.3 µM mRaichu, 30 µM GDP and 300 µM GTP. GAPd and GEFd were added to the cDT condition, as indicated. **: the probability of no difference between Ras activities under HP and AP was lower than the significance level (*t* = 0.01; Welch’s *t*-test).

We next prepared various conditions to examine how Ras activity responds to HP (Figure 2D, E). The concentrations of GDP and GTP in the cDT condition (0.3 µM mRaichu, 30 µM GDP, 300 µM GTP) were based on intracellular conditions^(43)^. When 10 µM GAPd or 1 µM GEFd was added to the cDT condition, the YFP/CFP ratio changed to levels similar to the GDP and GppCp condition, respectively (Figure 2D, E). Thus, in the cDT conditions, + 10 µM GAPd and 1 µM GEFd were judged to be saturated. Moreover, when both GAPd and GEFd were added (cDT + 1 µM GAPd + 0.1 µM GEFd), the YFP/CFP ratio reached a value between the saturation levels of GAPd and GEFd (cDT + 1 µM GAPd + 0.1 µM GEFd). Therefore, GAPd and GEFd can be used to regulate Ras activity. Ras activity [%], a measurement of the fraction of activated Ras under these conditions (see below for details), were calculated as shown in Table 1. The fraction of active Ras in normal cells was previously estimated as 20-40%^(44)^ and 30-70%, depending on the cell types^(45,46)^. Thus, the concentrations of GAPd and GEFd in the conditions of Table 1 were selected to reproduce the cellular Ras activities in normal and cancer cells.

**Table 1.** Summary of Ras activity [%] under various conditions.

| | +GAPd [μM] | +GEFd [nM] | Ras activity [%] | SD [%] | n |
| --- | --- | --- | --- | --- | --- |
| cDT | 1 | 1 | 29.1 | 6.6 | 6 |
| cDT | 1 | 10 | 32.6 | 0.6 | 6 |
| cDT | 1 | 100 | 36.8 | 11.4 | 6 |

The data were calculated from the YFP/CFP ratios measured under each condition. n: number of experiments.

### Establishment of *in vitro* system to measure HP sensitivity

Having characterized the response of mRaichu to varying concentrations of GTP, GDP, GAPd and GEFd, we next measured the temporal response of mRaichu to HP under those solution conditions. This was done using a newly established *in vitro* measurement system that included an HP-resistant optical cell consisting of inner and outer cells (Figure 3A). The mixture containing mRaichu and other components was brought to a quasi-static state by incubation for 1 h and then introduced into the inner cell, the space between the inner and outer cells was filled with deaerated water at 37 ℃, and the assembly was installed in the light path of fluorescence spectrophotometer B. HP was then indirectly applied to the reaction mixture in the inner cell by pressurizing the deaerated water between the inner and outer cells using a pump. Here, applied HP was 20-30 MPa, the higher end of the physiological range of HP on human chondrocytes^(38–40)^. Additionally, the effects of HP on the fluorescence properties of CFP and YFP moieties within mRaichu were tested by measuring the time-dependent change of fluorescence intensities at an excitation wavelength of CFP and YFP in the cDT + proteinase K condition with and without HP using the *in vitro* measurement system, respectively. Fluorescence intensity ratios in the cDT + proteinase K condition did not differ significantly over time and between the HP and AP conditions, so the effect of HP on the fluorescence properties of CFP and YFP moieties in mRaichu was negligible (Figure S4).

Changes of the YFP/CFP ratio of mRaichu were observed in the GDP, GppCp, cDT and cDT + proteinase K conditions over the time course of 20 min under AP and HP (Figure 3B). Rapid decreases of the YFP/CFP ratio were observed after HP was applied in all conditions except for the proteinase K-treated samples, but the degree of decrease was different among the conditions: the largest decrease was observed in the GppCp condition, followed by a smaller decrease in the cDT and GDP conditions (Figure 3B). Similar large decreases were observed when the concentration of GppCp was changed from 3 mM to 2 mM or 4 mM (Figure S5), indicating that the concentration of GppCp was saturated at 3 mM.

The possibility of a systematic error in the measurement system can be excluded because the decrease of the YFP/CFP ratio was negligibly small in the cDT + proteinase K condition. Here, the range of this decrease of mRaichu was smaller than that of Raichu (Figure S6), which indicated that the interaction between YFP and CFP moieties was partly responsible for the large HP-induced drop of the YFP/CFP ratio of the original Raichu.

Additionally, based on the result shown in Figure 3B, all subsequent observations were terminated at 10 min after the application of HP. We estimated that consumption of GTP by 0.3 µM mRaichu in 60-70 min, which includes the preincubation period and the observation period, is negligibly small compared with the original concentration of 30 µM in the cDT condition (for details, see Supplemental Information and Figure S7).

Unexpectedly, the YFP/CFP ratios of mRaichu in various identical solution conditions were significantly different when measured using the two fluorescence spectrometers: GDP, GppCp and cDT conditions and the cDT conditions containing various concentrations of GAPd and GEFd (Figure S8A). Similar levels of differences were observed even when the same normal quartz optical cell was used in the two fluorescence spectrometers (Figure S8A, B). Therefore, this difference should be due to the different wavelength dependence of the sensitivity of the photodetectors in the two fluorescence spectrometers. Thus, in subsequent experiments, the YFP/CFP ratios obtained using the HP-resistant cell and fluorescence spectrometer B were corrected to allow a direct comparison with the YFP/CFP ratios obtained using fluorescence spectrometer A in Figures 1 and 2 (for details, see Supplemental Information and Figure S8B).

Referring to the HP-induced YFP/CFP ratio decrease in Figure 3B, we introduced “Ras activity [%]”, which evaluates the YFP/CFP ratio under a given condition relative to that in the presence of GDP and in the presence of GppCp measured using the same optical cell (i.e., the conventional cell or the HP-resistant cell). More specifically, Ras activity [%] was calculated based on the YFP/CFP ratio as a linear interpolation between the ratio in the GDP condition set at 0% Ras activity [%] and that of the GppCp condition set at 100% Ras activity [%], in order to quantitatively evaluate the fraction of active Ras when the YFP/CFP ratio decreases by HP even when Ras should be fully active in the presence of 3 mM GppCp. Ras activities [%] in three basic conditions were calculated from the YFP/CFP ratios of mRaichu using the two measurement systems (normal quartz cell in fluorescence spectrophotometer A and HP-resistant cell in fluorescence spectrophotometer B) (Figure 3C). As expected, there were only small statistically insignificant differences between the Ras activity [%] values measured using the two systems under all conditions (t = 0.05 by Welch’s *t*-test). Hence, in the following experiments, Ras activity [%] has been used to evaluate the fraction of active Ras in mRaichu. Here, using the method of calculation of Ras activity [%], since the correction of the YFP/CFP ratio is a linear complement, the effect of the correction on Ras activity [%] is canceled.

**Figure 3.**
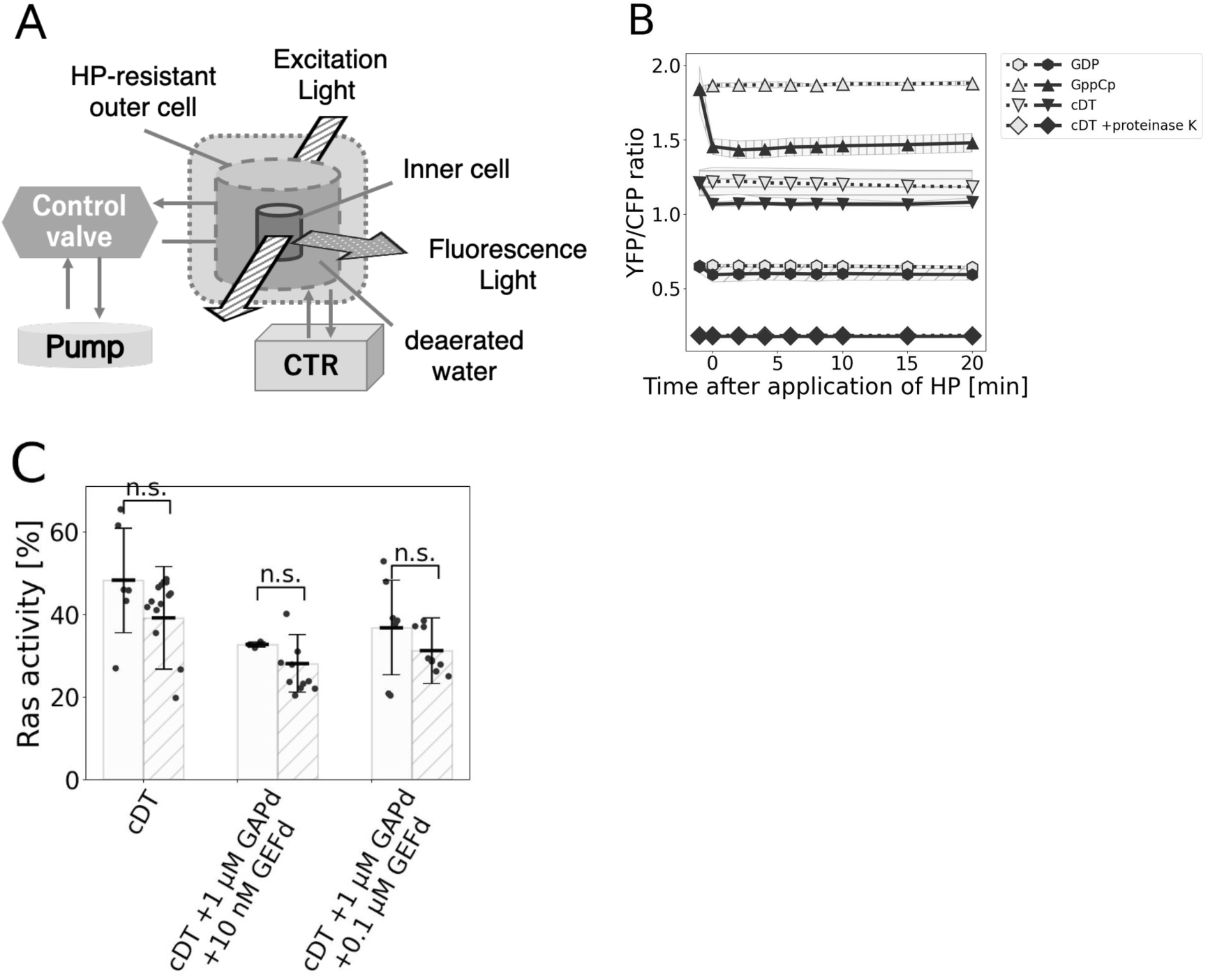
*In vitro* system to measure fluorescence spectra under HP and its method of analysis. (A) Overall configuration of the *in vitro* mRaichu activity measurement system under HP. CTR: constant temperature reservoir. (B) Time-dependent changes of the YFP/CFP ratio of mRaichu after application of HP (solid lines) and without HP application (AP; dotted lines) (average of 6 measurements). In this graph, as well as in all the following time course measurements, the values before the application of HP are shown at -1 min and the values when measurements began are shown at 0 min. The elapsed time from the onset of applying HP with the pump to obtaining the measured values was approximately 1 min. (C) Ras activity [%] ± SD calculated from the YFP/CFP ratios in the absence and presence of GAPd and GEFd (measured by fluorescence spectrometer A (left white bar of each group) and fluorescence spectrometer B (right hatched bar of each group) (average of 6 measurements each). The horizontal black bars and error bars indicate the average value ± SD. Excitation was at 433 nm.

### HP responsive factor(s) of the Ras-cycle

To identify the HP-responsive factor(s) within the Ras-cycle, we measured Ras activity [%] under HP in non-physiological conditions and compared each value with that measured under AP (Figure 4). These conditions were prepared to evaluate the HP sensitivity of mRaichu itself, mRaichu + GAPd and mRaichu + GEFd, respectively. There was a small but statistically significantly positive HP response of mRaichu itself (Figure 4A, B), which is consistent with the result of a prior *in silico* study^(31)^. In the case of mRaichu + GAPd, the difference showed responses depending on the GAPd concentration (Figure 4A, C, D, E). In the cDT condition containing a low GAPd concentration (10 nM), the HP-induced increase of Ras activity was larger than that in the absence of GAPd (Figure 4A, C). This HP-induced increase was also observed in the cDT condition containing another low GAPd concentration (100 nM). This increase was equivalent to that in the absence of GAPd, whereas the addition of a high concentration (10 µM) of GAPd extinguished the increase (Figure 4A, D, E). In contrast, in the case of mRaichu + GEFd, Ras activity [%] consistently increased by ∼27% within 1 min after the application of HP, and the GEFd-dependent increase was detectable even at the lowest GEF concentration tested (10 nM) (Figure 4A, F, G). The significance of these HP-induced increases was evaluated by comparing Ras activities with and without HP application at 5 min after HP application (Figure 4H and Table S1). Hence, the positive HP sensitivity resides in mRaichu itself and in the interaction between mRaichu and GAPd and GEFd. Between those three factors, in the presence of a low concentration of GAPd (10 nM), the mRaichu-GAPd interaction plays a major role (6.9%, assuming that the 16.3% increase includes contributions from the two mechanisms) in the HP response. In contrast, mRaichu-GEFd interaction also had large effects on the HP response of the Ras-cycle, independent of GEFd concentration (9.3-17.2%, assuming the same assumptions as for GAPd), equivalent to the mRaichu-GAPd interaction.

**Figure 4.**
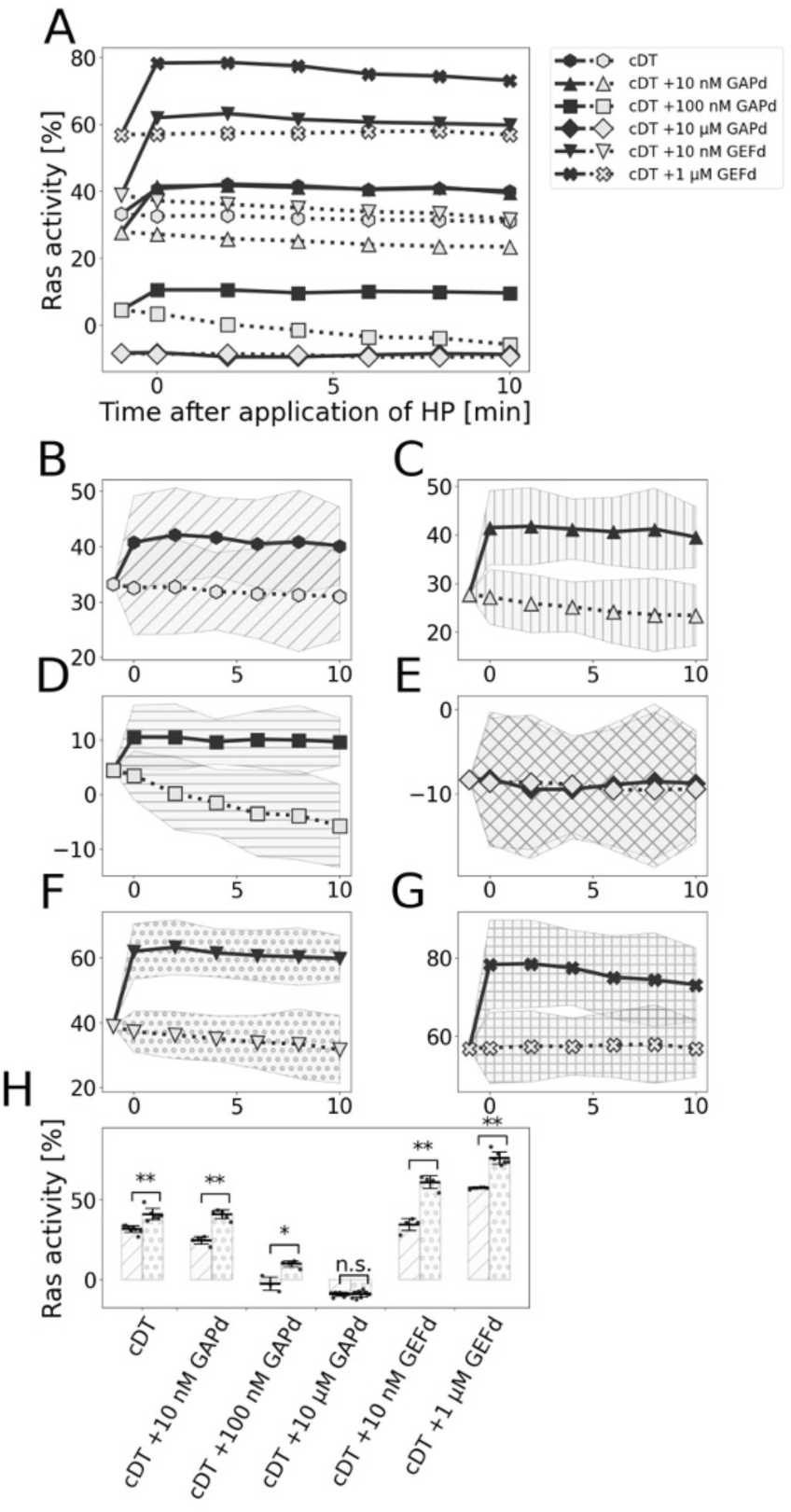
HP-response of Ras activity [%] in non-physiological conditions. (A-H) Time-dependent change of Ras activity [%] of mRaichu in the absence or presence of GAPd or GEFd after the application of HP (solid line) and without the application of HP (AP; dotted line) (average of 6-12 measurements each, as shown in Table S1). (B-G) show traces in (A) with an expanded Y-axis optimized for each trace. (H) Summary of Ras activity [%] differences under AP (left hatched bar of each group) and HP at 5 min after HP was applied (right dotted bar of each group). The horizontal black bars and error bars indicate the average value ± SD. *, **: the probability of no difference between Ras activities under HP and AP is lower than the significance level (*t* = 0.05, 0.01, respectively; Welch’s *t*-test).

### HP response in the presence of GAPd and GEFd

To obtain insight into the effect of the observed HP sensitivity of the Ras-cycle on cells, we measured Ras activity [%] under HP in the presence of both GAPd and GEFd and compared each value with that measured under AP (Figure 5 and Table S2). There were significant HP-induced increases in Ras activity [%] under the cDT conditions additionally containing GAPd and GEFd (Figure 5A, C, D, E, F, G and Table S2). Importantly, these results suggested that the Ras-cycle in cells has a significant HP sensitivity since positive HP responses were detected in a range of GAPd and GEF concentrations. The HP-induced increase of Ras activity [%] was statistically significant under all the conditions tested (Figure 5H and Table S2). HP-induced Ras activations were observed in various conditions when both concentrations of GAPd and GEFd varied (Figure S9). In particular, Ras activity [%] of the cDT + 10 nM GAPd + 10 nM GEFd condition showed the largest HP-induced increase, as expected.

**Figure 5.**
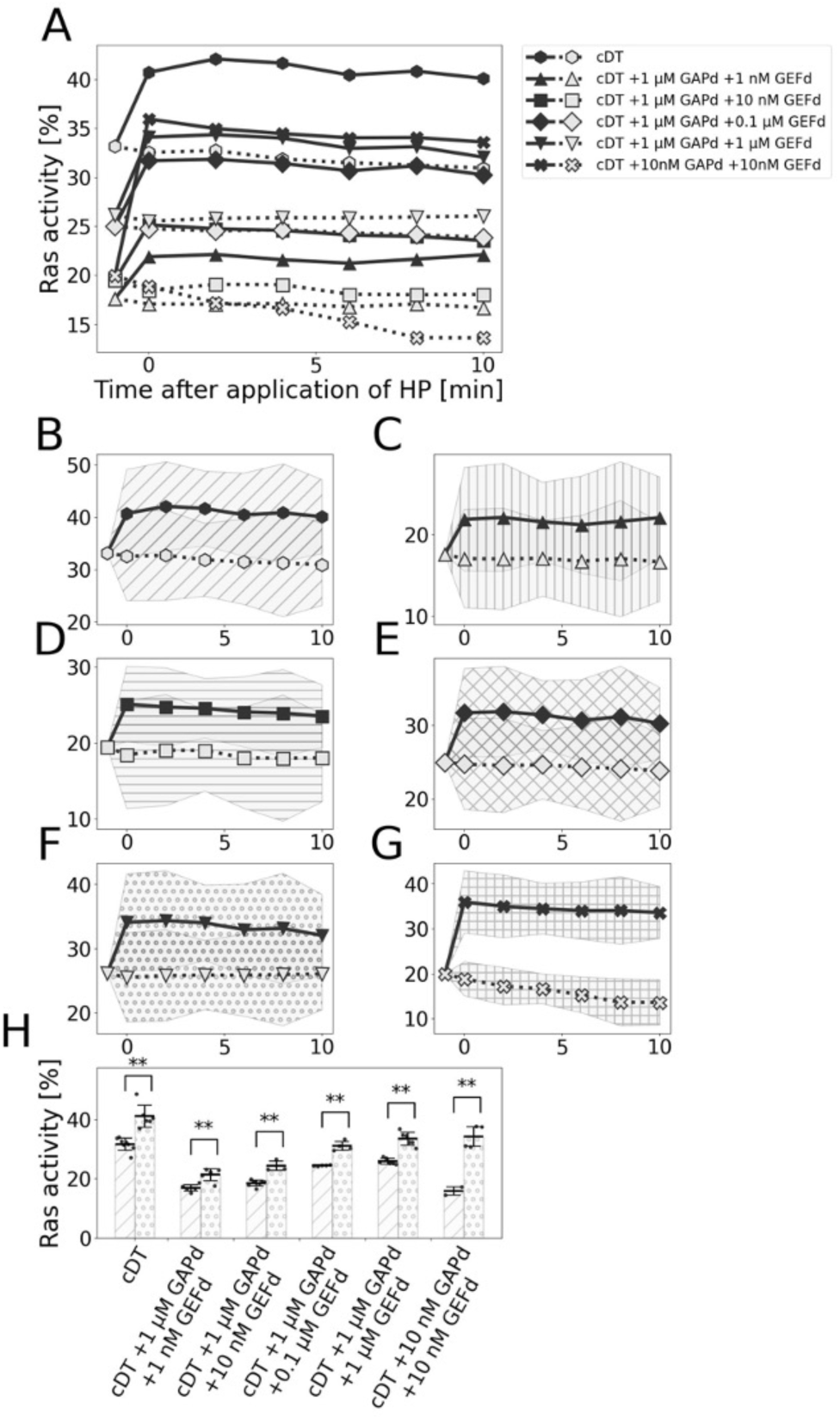
HP-response of Ras activity [%] in the co-presence of GAPd and GEFd. (A-H) Time-dependent change of Ras activity [%] of mRaichu in the absence or presence of GAPd or GEFd after the application of HP (solid line) and without the application of HP (AP; dotted line) (average of 3-9 measurements each, as shown in Table S2). (B-G) show traces in (A) with an expanded Y-axis optimized for each trace. (H) Summary of difference between Ras activity [%] under AP (left hatched bar of each group) and HP at 5 min after HP was applied (right dotted bar of each group). The horizontal black bars and error bars indicate the average value ± SD. **: the probability of no difference between Ras activities under HP and AP is lower than the significance level (*t* = 0.01; Welch’s *t*-test).

### Effect of GAPd on the observed HP sensitivity of the Ras-cycle

To evaluate the effect of GAPd concentration on the observed HP sensitivity in the co-presence of GAPd and GEFd, we examined the effects of HP on Ras activity [%] when GAPd concentration varied from the two normal cell mimic conditions: cDT + 1 µM GAPd + 1 nM GEFd and cDT + 1 µM GAPd + 10 nM GEFd (Figure 6 and Table S3). When GAPd concentration was decreased from 1 µM to 0.1 µM, comparable, significant levels of HP responses were observed (Figure 6A-C, F-H). In contrast, when GAPd concentration was increased to 10 µM, the HP sensitivity observed in the normal cell mimic condition became insignificant or weakened (Figure 6A, C, D, F, H, I). Figure 6E and J and Table S3 show that, at 5 min after HP was applied, Ras activity [%] clearly increased except when GAPd concentration was 10 µM.

**Figure 6.**
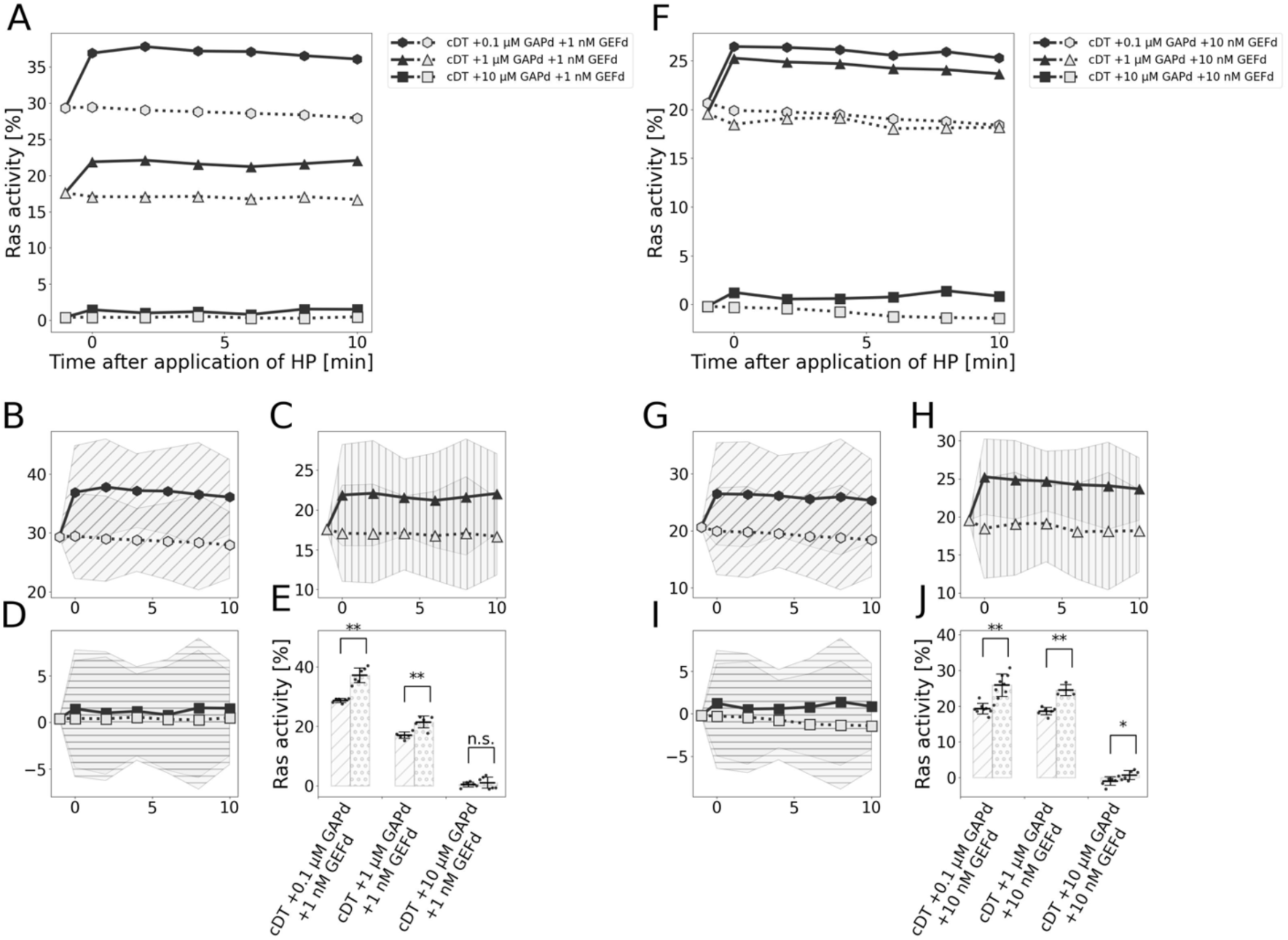
Effects of varying concentrations of GAPd on the HP response of the Ras-cycle in normal cell mimic conditions. (A-I) Time-dependent change of Ras activity [%] of mRaichu in the presence of various concentrations of GAPd and GEFd after the application of HP (solid line) and without the application of HP (AP; dotted line) (average of 3-9 measurements, each as shown in Table S3). (B-D) and (G-I) show traces in (A) or (F) with an expanded Y-axis optimized for each trace, respectively. (E, J) Summary of differences between Ras activity [%] under AP (left hatched bar of each group) and HP at 5 min after HP was applied (right dotted bar of each group). The horizontal black bars and error bars indicate the average value ± SD. **, *: the probability of no difference between Ras activities under HP and AP is lower than the significance level (*t* = 0.01, 0.05, respectively; Welch’s *t*-test).

Similarly, Figure 7 and Table S4 show Ras activity [%] under HP and AP when GAPd concentration was varied from the cancer cell mimic conditions (cDT + 1 µM GAPd + 0.1 µM GEFd). As was the case with the normal cell mimic conditions, a similar HP response was observed in the presence of 0.1 µM GAPd (Figure 7A, B, D). In the case of cancer cell mimic conditions, HP-induced Ras activation was significant, even in the presence of 10 µM GAPd (Figure 7A, C, D). Figure 7E and Table S4 show that, at 5 min after HP was applied, Ras activity [%]clearly increased in all the conditions tested, although the increase was much smaller when 10 µM GAPd was added. Hence, the positive HP sensitivities of the Ras-cycle were also decreased by excessive GAP in the cancer cell mimic condition.

**Figure 7.**
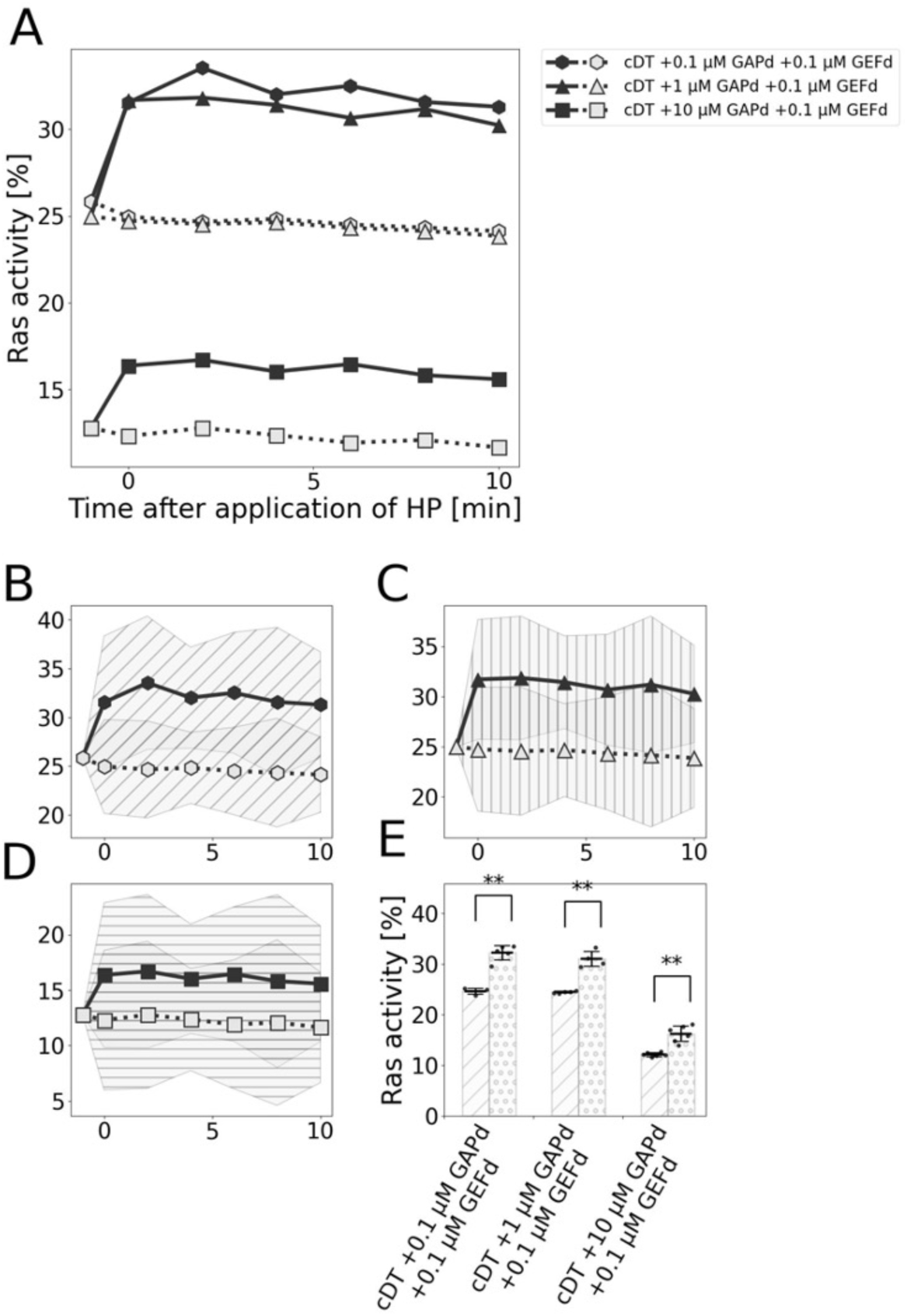
Effects of varying concentrations of GAPd on HP response of the Ras-cycle in cancer cell mimic conditions. (A-D) Time-dependent change of Ras activity [%] of mRaichu in the presence of various concentrations of GAPd and GEFd after the application of HP (solid line) and without the application of HP (AP; dotted line) (average of 5-7 measurements each, as shown in Table S4). (B-D) show traces in (A) with an expanded Y-axis optimized for each trace. (E) Summary of difference between Ras activity [%] under AP (left hatched bar of each group) and HP at 5 min after HP was applied (right dotted bar of each group). The horizontal black bars and error bars indicate the average value ± SD. **: the probability of no difference between Ras activities under HP and AP is lower than the significance level (*t* = 0.01; Welch’s *t*-test).

## Discussion

### Quantitative consideration of the YFP/CFP ratio and Ras activity [%]

We measured the YFP/CFP ratios of the fluorescence spectra of mRaichu when CFP was excited at 433 nm (Figure 1E, 2E). To compare the data in Figure 1E with those reported in a previous *in vivo* study on Raichu expressed in 293T cells, the YFP/CFP ratios were calculated from the fluorescence spectra of clarified cell lysate measured with the 433 nm excitation light in the previous study^(32,47)^ (Table 2). We conclude that YFP/CFP ratios of the present *in vitro* study and the previous *in vivo* study^(32)^, including the effects of GAPd/GAP and GEFd/GEF, are in reasonable agreement.

**Table 2.** Summary of the YFP/CFP ratio data in the previous *in vivo* study (Mochizuki et al. 2001) compared to that in this *in vitro* study.

| Source | Condition | YFP/CFP ratio ± SD |
| --- | --- | --- |
| <u>This study</u> | cDT | $1.25 \pm 0.13$ |
| | cDT + 10 μM GAPd | $0.89 \pm 0.02$ |
| | cDT + 1 μM GEFd | $1.96 \pm 0.05$ |
| | cDT + proteinase K | $0.05 \pm 0.00$ |
| <u>Mochizuki et al. (2001)</u> | wt (Raichu in 293T cells) | 1.6 |
|  | wt + GAP1m | 1.2 |
|  | wt + mSOS1 | 2.4 |
|  | wt + proteinase K | 0.5 |

Despite the A206K-equivalent point mutations introduced into CFP and YFP moieties in mRaichu, there were reproducibly significant and abrupt drops in the YFP/CFP ratio upon HP application in cDT, GDP and GppCp conditions. This could be due to the combination of a weak HP-sensitive interaction between inactive Ras and RafRBD and between active Ras and RafRBD. Regarding the HP-induced decrease of the YFP/CFP ratio in the presence of GppCp, we excluded the possibility that HP induced a dissociation of GppCp from Ras because the YFP/CFP ratio did not increase when the concentration of GppCp was increased from 2 mM to 4 mM (Figure S5).

To avoid the complexity due to this general HP-induced drop of the YFP/CFP ratios, we introduced a single common index, Ras activity [%]. With Ras activity [%], the YFP/CFP ratio of each measurement was evaluated relatively with respect to the ratio of the GDP condition set at 0% activity and the GppCp condition set at 100% in HP- and AP-experiments, respectively.

The data of this study and supplemental information are drawn from Figure 1E. wt, wt + GAP1m and wt + mSOS1: the YFP/CFP ratios of the previous study were measured from clarified lysates of 293T cells expressing Raichu or co-expressing Raichu and GAP1m or mSos1. wt + proteinase K: measured after the clarified cell lysate was treated with proteinase K.

### Identification of HP-sensitive elements in the Ras-cycle

We found that the activity ratio of mRaichu was weakly increased by HP (Figure 4 and Table S1). This HP sensitivity should reside in Ras and possible mechanisms for this are as follows: 1) HP weakens the interaction of Ras with GDP; 2) HP promotes the association of Ras to GTP; 3) HP inhibits the intrinsic GTPase activity of Ras. Among these possibilities, mechanism 3) is unlikely since NMR analysis of Ras itself showed no significant HP-induced structural changes in switch 1 and switch 2 regions^(31)^, implying that HP has little effect on the GTPase activity of Ras. Meanwhile, mechanisms 1) and 2) are qualitatively consistent with the result of *in silico* simulation in which the fraction of active Ras increased under HP^(31)^. Since the ligand-protein interaction is generally weakened by HP (reviewed by (20)), although there are some exceptions^(48,49)^, mechanism 2) is unlikely, leaving mechanism 1) as the most likely scenario.

We next evaluated the effect of GAPd and GEFd on HP-induced Ras activation (Figure 4 and Table S1). HP-induced activation was much more pronounced in the presence of 10 nM GAPd than in its absence (16.3 ±1.7% vs. 9.4 ±1.7%). Additionally, since the HP response increased significantly only when the GAPd concentration was low (10 nM) and based on the general tendency of protein complexes and protein-protein complexes to disassemble in HP (reviewed by (29)), we speculate that HP dissociates GAPd from the GAPd-Ras-GDP complex. Further assuming that HP accelerates dissociation of GDP from the resultant Ras-GDP complex, as discussed above, GTP would rapidly bind the apo Ras to generate active Ras, because GTP concentration is 10-fold higher than GDP in the cDT-based conditions and apo Ras has higher affinity for GTP than for GDP (reviewed by (20)). This would rapidly decrease the fraction of inactive Ras and increase that of active Ras.

In contrast, HP sensitivity of the Ras-cycle was observed in the presence of a wide range of GEFd concentrations (1-1000 nM). The mechanistic explanation for GEFd-enhanced HP sensitivity is more challenging than that in the presence of a low concentration of GAPd. However, based on the general tendency mentioned above (reviewed by (29)), the following mechanisms may contribute to the augmentation of HP-induced Ras activation by GEFd: 1) HP promotes the dissociation of GDP from the Ras-GDP-GEFd complex; 2) the intermediate state from GTP binding to the Ras-GEF complex to the GEF dissociation is over-stabilized by HP. Regarding mechanism 2), it is possible that, for example, Ras-GTP changes conformationally induced by HP and could assume a structure similar to that of Ras-GDP where the nucleotide is hardly exchanged because it is too stable^(37,50,51)^, despite GEF binding, but this mechanism contradicts the general tendency. Thus, we speculate that mechanism 1) is involved in the HP-induced Ras activation augmented by GEFd.

This HP-induced increase of Ras activity [%] was also observed in the co-presence of a varying combination of GAPd and GEFd concentrations, a condition mimicking cellular Ras activity (Figure 5, 6, 7 and Table S2, 3, 4). To further clarify the molecular mechanism of the HP sensitivity of the Ras-cycle in the physiological HP range, additional solution experiments, molecular dynamics (MD) simulations and other studies are needed.

### Physiological significance of the observed HP-induced Ras activation

Notably, the absolute Ras activity represented by the YFP/CFP ratio was reduced by HP under all the conditions tested (Figure 3B), and as discussed before, this may be due to the HP-induced partial dissociation of active Ras and RafRBD within mRaichu. Thus, in cells, even though HP increases the fraction of active Ras, downstream signaling through Raf kinase may decline, casting doubt on the physiological significance of our present finding. To address this concern, we speculate that in cellular environments there are additional mechanisms that stabilize the interaction between active Ras and Raf. For example, RafRBD in our mRaichu consists of a well-established minimum Ras Binding Domain of human Raf (reviewed by (52)). However, it was subsequently shown that cysteine-rich domain (CRD) downstream of RBD within Raf facilitates the binding of Raf-RBD to active Ras ^(53–55)^. Thus, inclusion of CRD to our mRaichu might stabilize active the Ras-RafRBD interaction under HP. The cytoplasm is a highly crowded environment and this may also stabilize the active Ras-RafRBD interaction under HP. These points should be investigated experimentally in the future.

The relative HP-induced increase of Ras activity (9.4 ± 1.7%) could have a meaningful physiological effect. Ras is involved in various cellular functions (reviewed by (12,13,19,20)) including cell growth^(24–26)^. The disruption of low-maintained Ras activity is a feature of cancer cells, and the relative increase of Ras activity by oncogenic mutations leads cells to a cancerous state (reviewed by (56)). Hence, this HP sensitivity of Ras may be involved in HP-induced modulation of various cellular functions of chondrocytes, and in certain cases, leads to oncogenesis. Particularly, in chondrocytes of mammalian tissues, a low level of HP stimulus (0.5-5 MPa) is not deleterious^(3–5)^ but high HP stimulus (25 MPa) render them pro-osteoarthritic^(6)^, leading to the development of cartilage osteosarcoma whose etiology is still unresolved (reviewed by (57)). Furthermore, because Ras has structural analogs such as Rho, Rap, etc. (reviewed by (20)) and the regulators of these proteins are structural analogs of GAPd or GEFd (reviewed by (28)), our result suggests that these proteins may also have HP sensitivity, similar to human Ras and its regulators. These possibilities can also be experimentally evaluated using a FRET-based probe similar to mRaichu. Of note, similar FRET-based probes for Rap1^(32)^ and Rho^(47)^ have already been constructed.

Taking a broader view of the natural world, organisms such as deep-sea fishes are constantly exposed to HP and whales that travel between the sea surface and deep sea are temporarily exposed to high HP. If the observed HP sensitivity is a general feature of the Ras-cycle in all vertebrates, deep sea fishes and whales might have acquired a system to prevent the unwanted HP-induced Ras activation. Alternatively, if the observed HP sensitivity of the human Ras-cycle is specific to organisms in which HP sensitivity has physiological significance, such as humans and other large land animals, it is possible that such HP sensitivity of the Ras-cycle has evolved from the non-HP-sensitive ancestral Ras system. Therefore, it would be interesting to examine the Ras-cycle of representative vertebrates including deep-sea fishes and whales in similar experiments.

## Author Contributions

T.M., K.F., T.U. and T.Q.P.U. conceived the present experiments, based on preliminary experiments performed by T.U. T.M. prepared the proteins. M.C. built the experimental circuit. T.M. carried out all experiments and analyzed the data. T.M. wrote the manuscript draft and T.Q.P.U. revised it. All authors discussed the results and commented on the manuscript.

## Acknowledgments

We are grateful to Prof. Michiyuki Matsuda of Kyoto University for generously providing the plasmid to express Raichu in human cells. This work was supported by Grants-in-aid from the Ministry of Education, Culture, Sports, Science and Technology to TU (No. 19H01173), and a fellowship from the JST-Mirai Program, Grant Number JPMJSP2128, Waseda University, and the Oshima scholarship foundation, Japan, to TM.

## Notes

### Competing Interest Statement

The authors have declared no competing interest.

