## Supplemental Materials for "Ras Activation by Hydrostatic Pressure is Enhanced by GAP and GEF *in vitro*"

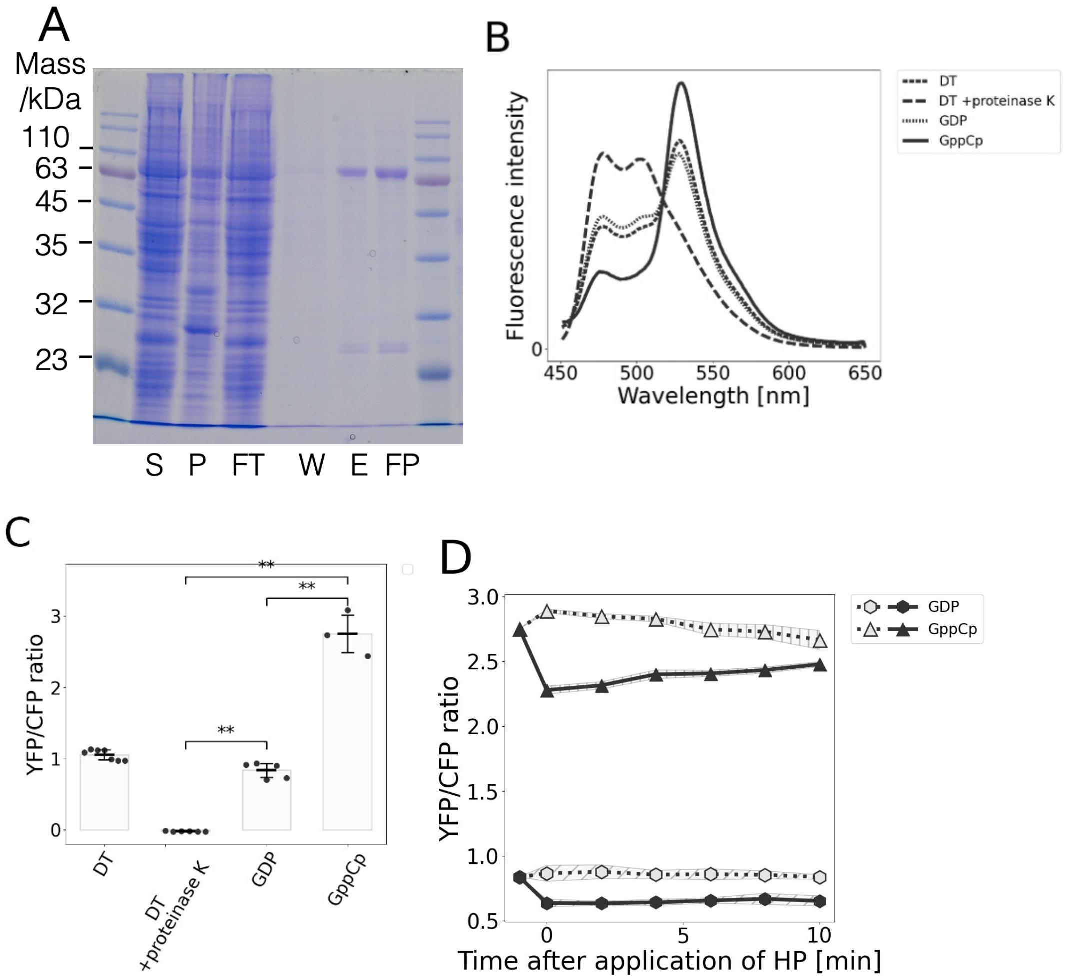


Figure S1. Purification of Raichu

(A) Purification of Raichu (predicted molecular mass is 87,000). S, P: Supernatant and pellet fractions after insect cell lysis and centrifugation, FT: Flow-through after incubation with Ni-NTA resin, W: Effluent with wash buffer, E: Eluted solution and FP: Final product after concentration and ultracentrifugation. (B) Difference in fluorescence spectra of Raichu depending on the solution conditions (average of 3 measurements each). Fluorescence spectra of the mixtures were measured at an excitation wavelength of 433 nm. (C) YFP/CFP ratio calculated from the data shown in (B). The horizontal black bars and error bars indicate the average value ± SD. GDP and GppCp conditions are the solution containing 0.3 μM Raichu and 1 mM GDP and 3 mM GppCp, respectively. DT and DT + proteinase K conditions are the solution containing 0.3 μM Raichu, 19.8 μM GDP and 13.2 μM GTP with or without 100 μg/mL proteinase K, respectively. (D) Time-dependent change of the YFP/CFP ratio of Raichu after the application of HP (solid lines) and without the application of HP (AP; dotted lines) (average of 6 measurements each). In this and all the following graphs showing the time course of YFP/CFP ratios after the application of HP, the values before HP application are shown at -1 min and the values when measurements began are shown at 0 min. The elapsed time from the onset of applying HP with the pump to obtaining the measured values was approximately 1 min. The YFP/CFP ratios in (D) were corrected in a manner explained in Supplemental Information, to allow for a direct comparison with the YFP/CFP ratios in (C).

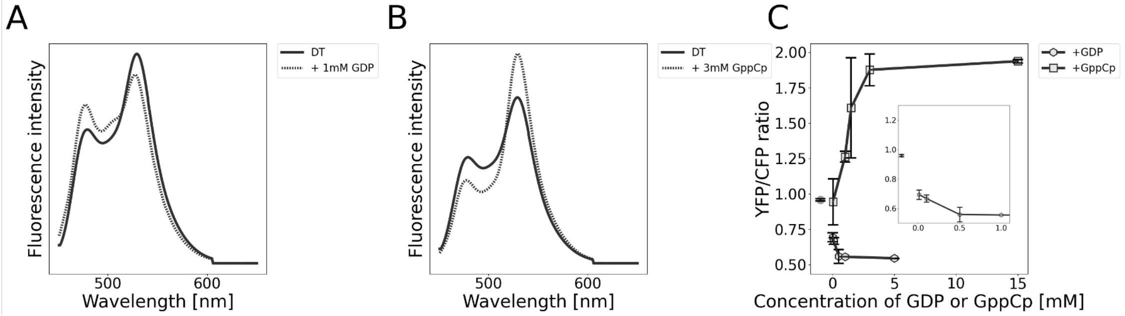


Figure S2. Effects of GDP and GppCp on the fluorescence spectra of mRaichu

(A, B) Fluorescence spectra of mRaichu in the presence of the 1 mM GDP (A) and 1 mM GppCp (B) (average of 3 measurements each). Fluorescence spectra of the mixtures were measured at an excitation wavelength of 433 nm. (C) The YFP/CFP ratio of mRaichu in the presence of various concentrations of GDP or GppCp (average ± SD of 3 independent measurements). The YFP/CFP ratio did not change significantly when GDP and GTP concentrations were increased above 1 mM, indicating that the effects of GDP and GTP are saturated at 1 mM for 0.3 μM mRaichu in the solution. The inset in (C) shows the enlarged view at low concentrations of GDP.

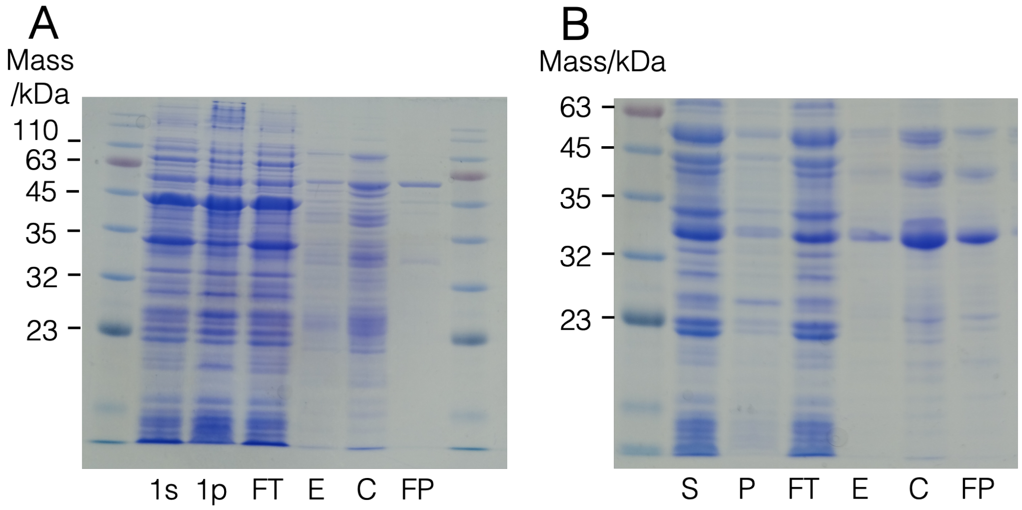


Figure S3. Purification of GAPd and GEFd.

(A, B) Purification of GEFd (A) (predicted molecular mass is 54,400) and GAPd (B) (predicted molecular mass is 26,600). 1s, 1p: Supernatant and pellet fractions after *E. coli* cell lysis and centrifugation, FT: Flow-through after incubation with Ni-NTA resin, E: Eluted solution, C: product after concentration, and FP: product after gel filtration chromatography (A) or anion exchange chromatography (B).

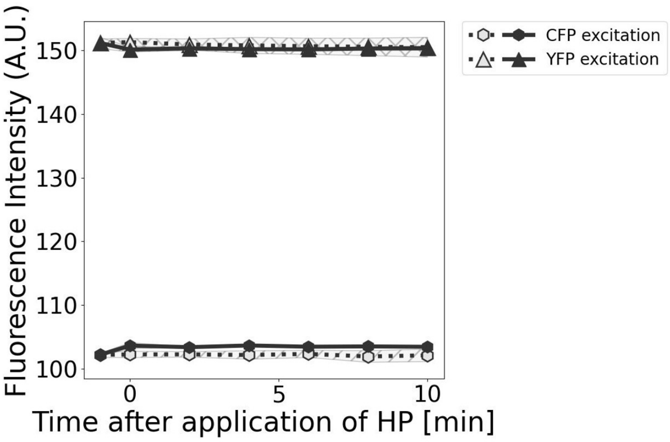


Figure S4. HP response attributed to CFP and YFP moieties.

mRaichu in cDT was treated with 100 μg/mL proteinase K, and time-dependent changes of the YFP and CFP fluorescence intensity (A.U.) after the application of HP (solid lines) and without the application of HP (AP; dotted lines) were measured. In the mRaichu solution treated with proteinase K, no significant change in fluorescence was observed in YFP and CFP fluorescence after HP was applied. Therefore, the possibility that fluorescent properties of CFP and YFP are affected by HP was eliminated.

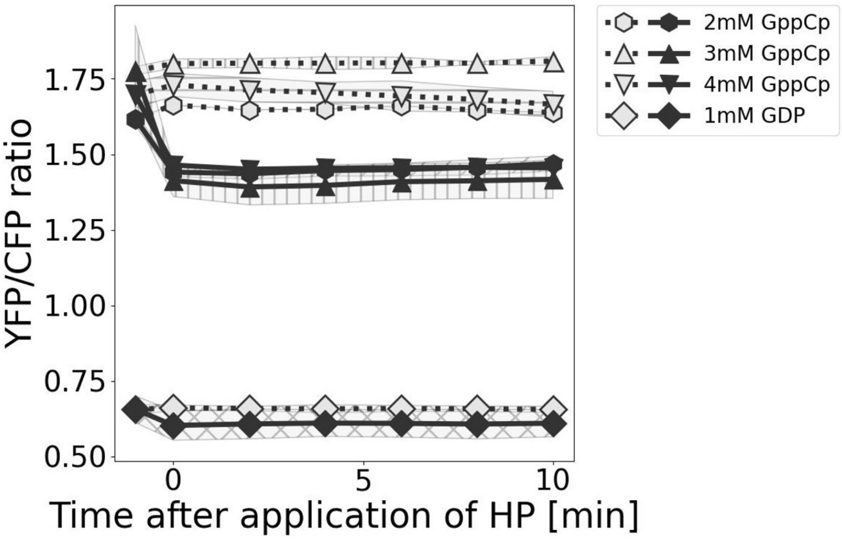


Figure S5. HP-induced spectral changes in the presence of various concentrations of GppCp.

Time-dependent changes of the YFP/CFP ratio after the application of HP (solid lines) and without the application of HP (AP; dotted lines) (average of 3 or 5 measurements). These conditions contained 0.3 μM mRaichu and 2, 3 or 4 mM GppCp, as indicated. The data in the GDP condition in Figure 3B is shown as the reference. The HP-induced decrease of YFP/CFP did not differ much when the concentration of GppCp was changed from 2 mM to 3 and 4 mM. Thus, we assumed this decrease is predominantly caused by the HP sensitivity of the interaction between active Ras and RafRBD, rather than due to the HP-sensitive binding of GppCp to apo-Ras.

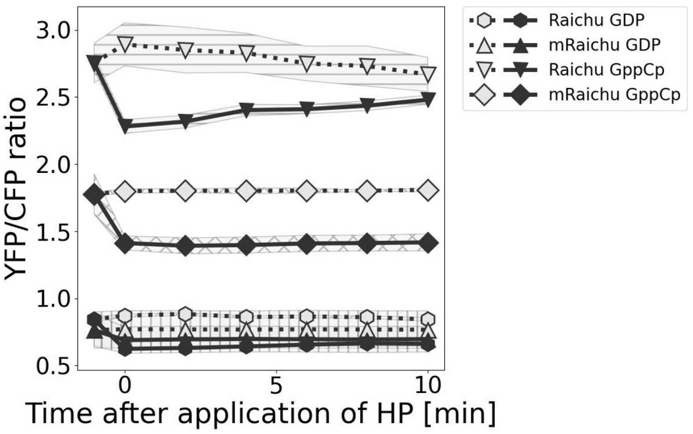


Figure S6. Effects of A206K mutations in YFP and CFP moieties within Raichu
Time-dependent changes of the YFP/CFP ratio after the application of HP (solid lines) and without the application of HP (AP; dotted lines) (average of 3 measurements each). The reaction solutions contained 0.3 µM Raichu or mRaichu and 1 mM GDP or 3 mM GppCp, as indicated. Immediately after HP was applied (-1-0 min), the YFP/CFP ratio decreased significantly by 0.24 ± 0.03 (mean ± SD) in the Raichu GDP condition. This HP-induced reduction of the YFP/CFP ratio should include an HP-induced dissociation of the YFP-CFP interaction. With mRaichu, the YFP/CFP ratio was slightly smaller than that of Raichu (Raichu: 0.84 ± 0.02 vs. mRaichu: 0.76 ± 0.24) and the HP-induced decrease was much smaller (Raichu: -0.24 ± 0.03 vs. mRaichu -0.08 ± 0.02; t < 0.05; Welch's *t*-test). Additionally, comparing the change of the YFP/CFP ratio in the presence of GppCp, the absolute value of the YFP/CFP ratio was smaller (Raichu: 2.75 ± 0.20 vs. mRaichu: 1.78 ± 0.09), as was the HP-induced decrease (Raichu: -0.50 ± 0.21 vs. mRaichu: 0.28 ± 0.09) by introducing the mutations. These results suggest that introducing the A206K equivalent mutations within Raichu disrupts the YFP-CFP interaction, whereas the weak HP-sensitive interaction between inactive Ras and RafRBD within mRaichu remains.

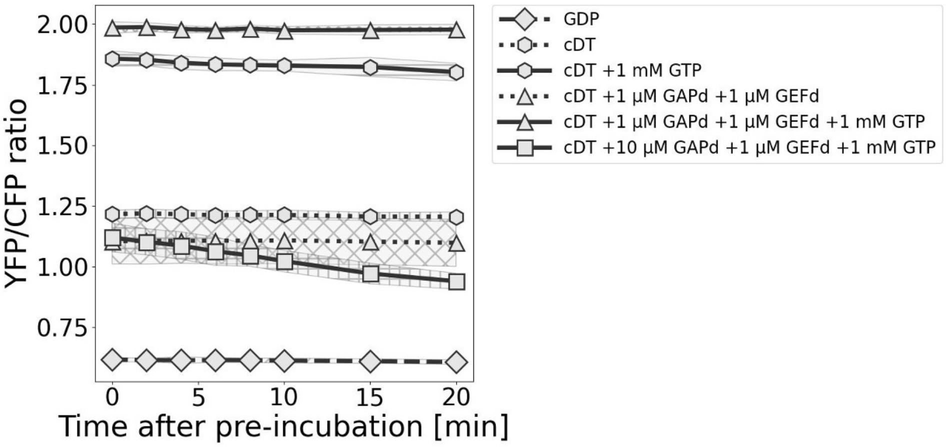


Figure S7. Temporal change of the YFP/CFP ratio under ambient pressure in the presence of an excess amount of GTP

Time-dependent change of the YFP/CFP ratio of mRaichu in the cDT condition and the cDT conditions additionally contained GAPd, GEFd and GTP, as indicated (average of 3 or 6 measurements each). The values when measurements began are shown at 0 min after 1 h of solution preparation and pre-incubation at 37 °C. Since the YFP/CFP ratios change depending on the GTP concentration and these ratios are always higher than that in the GDP condition, at least, the GTP in solution is not depleted (Figure 2E). Likewise, at least, in all cDT-based conditions in Figure S7, GTP in solution is not depleted. Additionally, by checking GTP depletion over time, the YFP/CFP ratios of all the cDT-based conditions, except for the cDT and cDT + 1 μM GAPd + 1 μM GEFd condition, which has a very high GTPase activity, did not change temporally during the 20 min incubation, indicating that the GTP concentrations were kept constant during this period. This is consistent with the result of kinetic simulation (for details, see Supplemental Information), indicating that the solution after 1 h of pre-incubation reached a quasi-equilibrium state. However, in the cDT + 10 μM GAPd + 1 μM GEFd + 1 mM GTP condition, which was not used in this study to analyze the HP-induced response of mRaichu, a linear decrease of the YFP/CFP ratio was detected, reflecting a high Ras-GTPase activity due to high concentrations of GAPd, GEFd and GTP.

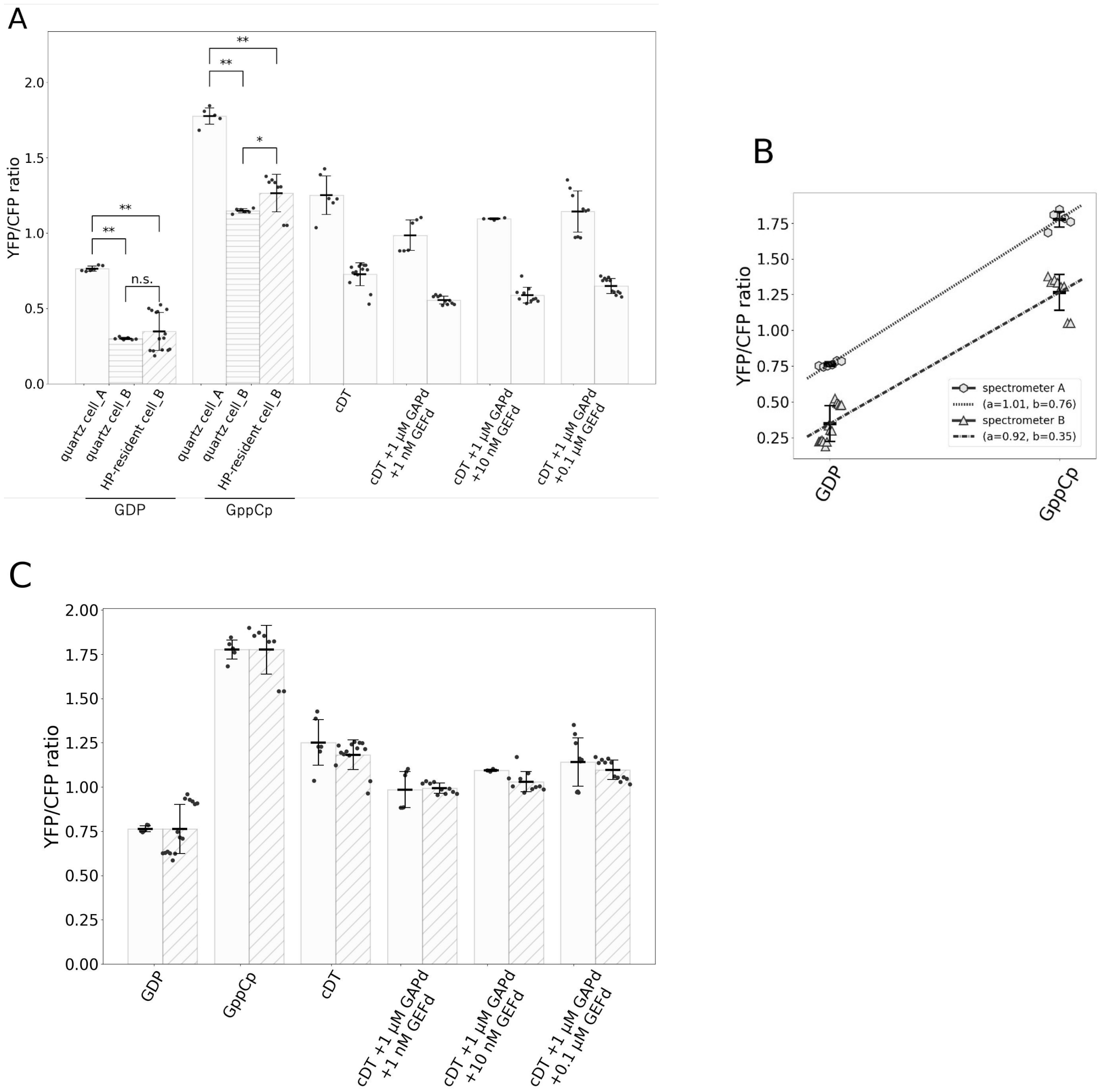


Figure S8. The difference of the *in vitro* experimental system.

(A) YFP/CFP ratios of mRaichu in various conditions were measured with a conventional quartz cell set in the fluorescence spectrometer A (left white bar of each group) and the HP-resistant cell set in the fluorescence spectrometer B (right hatched bar of each group). The data were taken from Figure 2E and 3B, respectively. Additionally, in GDP and GppCp conditions, YFP/CFP were also obtained using the quartz cell in the fluorescence spectrometer B (average of 6 measurements) (middle bar with horizontal lines of the group). '_A' and '_B' indicate that the data were obtained using fluorescence spectrometers A and B, respectively. Excitation was at 433 nm. GDP and GppCp conditions are the solutions containing 0.3 μM mRaichu, 1 mM GDP or 3 mM GppCp, respectively. The cDT-based conditions contained 30 μM GDP and 300 μM GTP with various concentrations of GAPd and GEFd, as indicated. In all solution conditions, the YFP/CFP ratio measured with the HP-resistant optical cell in fluorescence spectrometer B was smaller than that measured with the conventional optical cell in fluorescence spectrometer A. Additionally, the YFP/CFP ratio is predominantly changed by differences in the fluorescence spectrometer rather than the cell type (normal quartz cells vs. HP-resistant cells). (B) YFP/CFP ratios of mRaichu derived from the data obtained using the quartz cell in the fluorescence spectrometers A and the data in the HP-experiment (i.e., the HP-resistant cell in the fluorescence spectrometers B) shown in (A). a, b: the slope and y-intercept of the linear lines. These lines were obtained by plotting each data point with the YFP/CFP ratios as y-values and x-value of the GDP condition set at 0 and the GppCp condition at 1, respectively, and the slope and the y-intercept of the lines are shown as a and b in (B), respectively. To allow quantitative comparison of data obtained using the two fluorescence spectrophotometers, this was corrected by linear transformation to the YFP/CFP ratio obtained in the HP experiment. For details, see Supplemental Information. (C) Corrected YFP/CFP ratios of the data in the HP-experiment in (A). This correction significantly reduced the differences of the YFP/CFP ratios due to the differences in fluorescence spectrophotometers. In all graphs, the horizontal black bars and error bars indicate the average value ± SD.

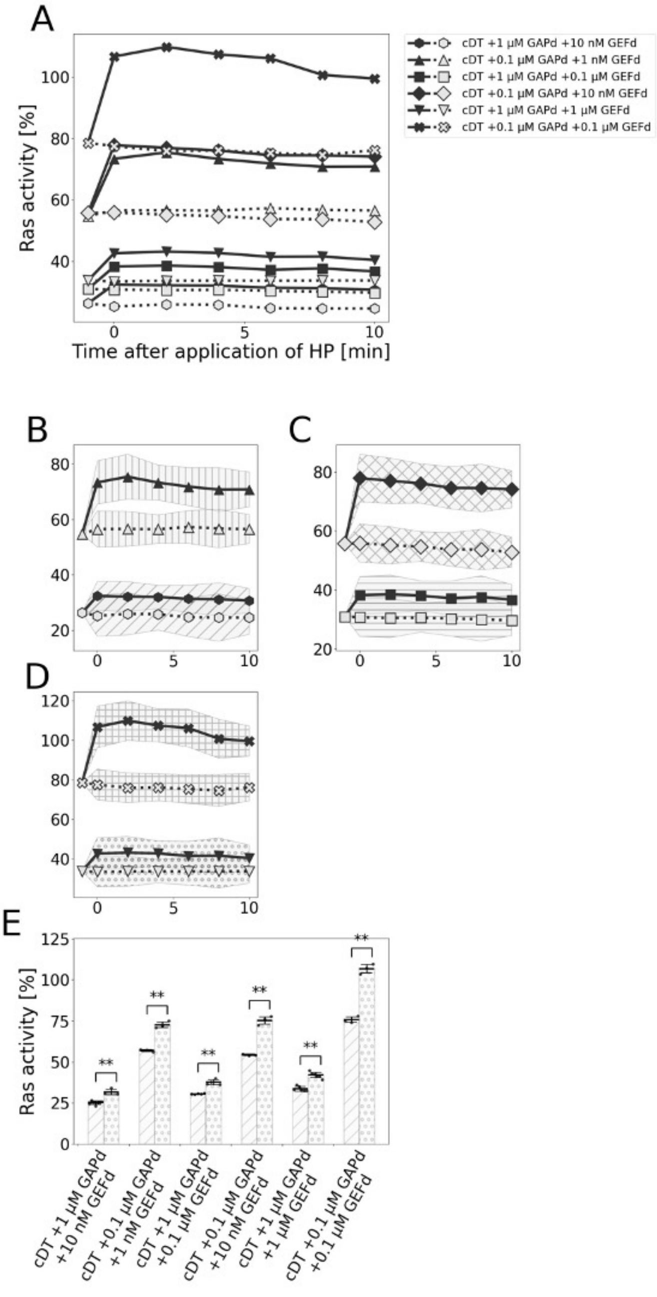


Figure S9. HP-response of Ras activity in the presence of various concentrations of GAPd and GEFd.

(A-D) Time-dependent change of Ras activity [%] of mRaichu in the presence of GAPd and GEFd after the application of HP (solid line) and without the application of HP (AP; dotted line) (average of 6 measurements each). (B-D) show traces in (A) with an expanded Y-axis optimized for each trace. (E) Summary of Ras activity differences under AP (left hatched bar of each group) and HP at 5 min after HP was applied (right dotted bar of each group). The horizontal black bars and error bars are the average value ± SD. (B)-(E) show the results of paired conditions where both GAPd and GEFd concentrations in one condition were reduced to one-tenth in the other. Notably, in all paired conditions, HP-induced Ras activations were greater when concentrations of both GAPd and GEFd were reduced by one-tenth, as summarized in Table S5.

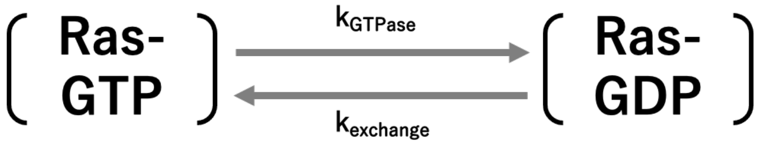


Figure S10. Simple kinetic model of the Ras-cycle.

The reaction rate ($k$) for each is as follows. $k_{GTPase}$: GTP hydrolysis rate in GTP-bound Ras (GTP-Ras) ($5.1 \times 10^{-4}$ [$s^{-1}$] (at 37 °C)[(58)]). $k_{exchange}$: The exchange rate of GDP on Ras to GTP ($1.8 \pm 0.2 \times 10^{-4}$ [$s^{-1}$] (at RT)[(59)]). In this model, since GDP binding rate to apo-Ras is very small compared to the GDP dissociation rate to form apo-Ras in the presence of an intracellular concentration of GDP, this reaction was ignored (reviewed by (20), (23)). Similarly, the reverse reaction of GTP hydrolysis from GDP-Pi-Ras to GTP-Ras is negligibly small[(58)]. $k_{GTPase}$ was set to $0.8 \pm 0.1$ [$s^{-1}$] in the presence of 1 μM GAP (at 37 °C[(60, 61)], and $k_{exchange}$ was set to $5.0 \pm 1.0 \times 10^{-3}$ [$s^{-1}$] in the presence of 1 μM GEF (at RT)[(59)]. These two reaction rates were assumed to be maximum constant values because YFP/CFP ratios were saturated to the upper or lower limit after adding 1 μM GAPd or GEFd, respectively (Figure 2C).

Table S1. Summary of HP-response of Ras activity [%] in non-physiological conditions

| | **+GAPd [μM]** | **+GEFd [nM]** | **Average change [%]** | **SD [%]** | **n (AP)** | **n (HP)** |
|---|---|---|---|---|---|---|
| **cDT** | - | - | 9.4** | 1.7 | 8 | 6 |
| **cDT** | 0.01 | - | 16.3** | 1.7 | 4 | 5 |
| **cDT** | 0.1 | - | 12.3* | 2.6 | 3 | 4 |
| **cDT** | 10 | - | -0.0 | 0.8 | 9 | 9 |
| **cDT** | - | 0.01 | 26.6** | 2.7 | 4 | 4 |
| **cDT** | - | 1 | 18.7** | 1.6 | 6 | 6 |

The data in Figure 4H are shown. “n”: number of HP and AP experiments. *, **: probability of no difference between Ras activities under HP and AP is lower than the significance level ($t$ = 0.05, 0.01, respectively; Welch’s $t$-test) .

Table S2. Summary of HP-response of Ras activity [%] in the co-presence of GAPd and GEFd

| | **+GAPd [μM]** | **+GEFd [nM]** | **Average change [%]** | **SD [%]** | **n (AP)** | **n (HP)** |
|---|---|---|---|---|---|---|
| **cDT** | - | - | 9.4** | 1.7 | 8 | 6 |
| **cDT** | 1 | 1 | 4.5** | 1.0 | 5 | 5 |
| **cDT** | 1 | 10 | 5.8** | 1.0 | 7 | 3 |
| **cDT** | 1 | 100 | 6.6** | 0.7 | 5 | 4 |
| **cDT** | 1 | 1000 | 7.6** | 1.0 | 6 | 6 |
| **cDT** | 0.01 | 10 | 18.3** | 2.0 | 2 | 4 |

The data in Figure 5H are shown. "n": number of HP and AP experiments. **: probability of no difference between Ras activities under HP and AP is lower than the significance level ($t$ = 0.01; Welch's $t$-test).

Table S3. Summary of effects of varying concentrations of GAPd on the HP response of Ras-cycle in normal cell mimic conditions

| | **+GAPd [µM]** | **+GEFd [nM]** | **Average change [%]** | **SD [%]** | **n (AP)** | **n (HP)** |
|---|---|---|---|---|---|---|
| **cDT** | 0.1 | 1 | 8.5** | 1.0 | 6 | 6 |
| **cDT** | 1 | 1 | 4.5** | 1.0 | 5 | 5 |
| **cDT** | 10 | 1 | 0.6 | 0.8 | 6 | 6 |
| **cDT** | 0.1 | 10 | 6.6** | 1.2 | 8 | 9 |
| **cDT** | 1 | 10 | 5.9** | 1.0 | 7 | 3 |
| **cDT** | 10 | 10 | 1.7* | 0.7 | 6 | 6 |

The data in Figure 6E and 6J are shown. The first and fourth lines show the two normal cell mimic conditions and the increased concentration of GAPd from those conditions. “n”: number of HP and AP experiments. **, *: probability of no difference between Ras activities under HP and AP is lower than the significance level ($t$ = 0.01, 0.05, respectively; Welch’s $t$-test).

Table S4. Summary of effects of varying concentrations of GAPd on HP response of Ras-cycle in cancer cell mimic conditions

| | **+GAPd [μM]** | **+GEFd [nM]** | **Average change [%]** | **SD [%]** | **n (AP)** | **n (HP)** |
|---|---|---|---|---|---|---|
| **cDT** | 0.1 | 100 | 7.6** | 0.7 | 3 | 5 |
| **cDT** | 1 | 100 | 6.6** | 0.7 | 5 | 4 |
| **cDT** | 10 | 100 | 4.1** | 0.6 | 6 | 6 |

The data in Figure 7E are shown. The first line shows the cancer cell mimic condition and the increased concentration of GAPd from that condition. "n": number of HP and AP experiments. **: probability of no difference between Ras activities under HP and AP is lower than the significance level ($t$ = 0.01; Welch's $t$-test).

Table S5. Summary of HP-response of Ras activity [%] in conditions under which the ratio of GAPd to GEFd concentrations were kept constant.

| | +GAPd [μM] | +GEFd [nM] | Average change [%] | SD [%] | n (AP) | n (HP) |
|---|---|---|---|---|---|---|
| **cDT** | 1 | 10 | 6.5** | 1.0 | 7 | 3 |
| **cDT** | 0.1 | 1 | 15.7** | 1.1 | 3 | 3 |
| **cDT** | 1 | 100 | 7.1** | 0.8 | 5 | 4 |
| **cDT** | 0.1 | 10 | 21.1** | 1.3 | 3 | 3 |
| **cDT** | 1 | 1000 | 8.4** | 0.9 | 6 | 6 |
| **cDT** | 0.1 | 100 | 31.1** | 1.7 | 3 | 3 |

The data in Figure S9E are shown. Note that the concentration ratio of GAPd/GEFd is the same for lines 1 and 2, lines 3 and 4 and lines 5 and 6. "n": number of HP and AP experiments. **: probability of no difference between Ras activities under HP and AP is lower than the significance level ($t = 0.01$; Welch's *t*-test).

### Supplemental Information

Correction method of the YFP/CFP ratio obtained in the HP experiment

The slopes and y-intercepts of the two lines connecting the YFP/CFP ratios in the GDP and GppCp conditions (Figure S8B) were different. Thus, the values of the YFP/CFP ratio differ depending on the fluorescence spectrometer even though Ras activity under the same solution conditions must be the same regardless of the fluorescence spectrophotometer. Therefore, the YFP/CFP ratios measured by spectrometer B were corrected by a linear transformation ($\alpha x + \beta$, $\alpha$: 1.10, $\beta$: 0.38, x: the YFP/CFP ratio) to be consistent with that measured by spectrophotometer A.

GTP concentration change in experimental solutions during the time scale of this experiment system

The change in GTP concentration during the measurement was calculated using a simple kinetic model (Figure S10). For simplicity, we hypothesized that the transitions between the inactive and active states of Ras occur with the given reaction rates. In our current series of measurements, the fastest consumption of GTP should occur in the cDT + 10 μM GAPd + 1 μM GEFd condition. Based on these assumptions, the maximum change of GTP concentration was 8.9 μM in the timescale of this measurement, 70 min (preincubation time: 60 min, measurement time: 10 min) and the changes of GTP and GDP concentrations were calculated as Table S6 during the 10 min of the measurement. These changes were negligibly small for the total initial concentrations (GTP: 300 μM and GDP: 30 μM). Therefore, we concluded that the solution was in a quasi-stable state within this measurement time and that the change in GTP/GDP concentrations did not significantly contribute to the fluorescence spectral change of mRaichu, although the starting condition (condition when we started the measurement) had somewhat drifted from the initial condition. These estimates are consistent with the fact that the YFP/CFP ratio did not change significantly during the 10 min incubation (Figure S7).

Table S6. Summary of the estimated change of GTP concentration

| | | Starting concentration [μM] | Final concentration [μM] |
|---|---|---|---|
| cDT condition | GTP concentration | 298.4 | 298.1 |
| | GDP concentration | 31.6 | 31.9 |
| cDT + 10 μM GAPd +1 μM GEFd condition | GTP concentration | 246.6 | 237.7 |
| | GDP concentration | 83.4 | 92.3 |

These values of GTP and GDP concentrations in cDT and cDT + 10 μM GAPd + 1 μM GEFd conditions were calculated using the characteristic reaction rates (Figure S10), respectively. Starting and final concentrations indicate concentrations at the start of measurement after pre-incubating the solution for 60 min at 37 °C and at the end of the 10 min measurement, respectively. The difference between the initial concentrations (GTP: 300 μM and GDP: 30 μM) and the starting concentrations at the beginning of the measurement, and the difference between the starting and final concentrations during the measurement, are negligibly small compared with the initial concentration.
